# Impact of bioactive molecule inclusion in lyophilized silk scaffolds varies between *in vivo* and *in vitro* assessments

**DOI:** 10.1101/2022.05.24.493207

**Authors:** Julie F Jameson, Marisa O Pacheco, Elizabeth C Bender, Nisha M Kotta, Lauren D Black, David L Kaplan, Jonathan M Grasman, Whitney L Stoppel

## Abstract

Biomaterials can influence the coordinated efforts required to achieve tissue rehabilitation. Sponge-like silk fibroin scaffolds that include bioactive molecules have been shown to influence tissue repair. However, the mechanisms by which scaffold formulations elicit desired *in vivo* responses is unclear. Here, acellular silk scaffolds consisting of type I collagen, heparin, and/or vascular endothelial growth factor (VEGF) were used to investigate material fabrication and composition parameters that drive scaffold degradation, cell infiltration, and adipose tissue deposition *in vivo*. In subcutaneous implants, scaffold degradation was assessed, and results show that the percentage of cells infiltrating the scaffold increased when scaffold formulations contained bioactive molecules. To gain further insight, calculated *in vitro* enzymatic degradation rates increased with higher enzyme concentrations and theoretical cleavage sites. However, the addition of type I collagen and heparin to the scaffold at relevant concentrations did not change degradation rates, compared to silk alone. These *in vitro* results are contrary to observations *in vivo*, where bioactive molecules influence local protein deposition, immune cell infiltration rates, and vascularization. Thus, quantitative *in vitro* and *in vivo* evaluations aid in determining the mechanisms by which biomaterials influence tissue repair and support intentional biomaterial design for clinical applications.

**Graphical Abstract:** 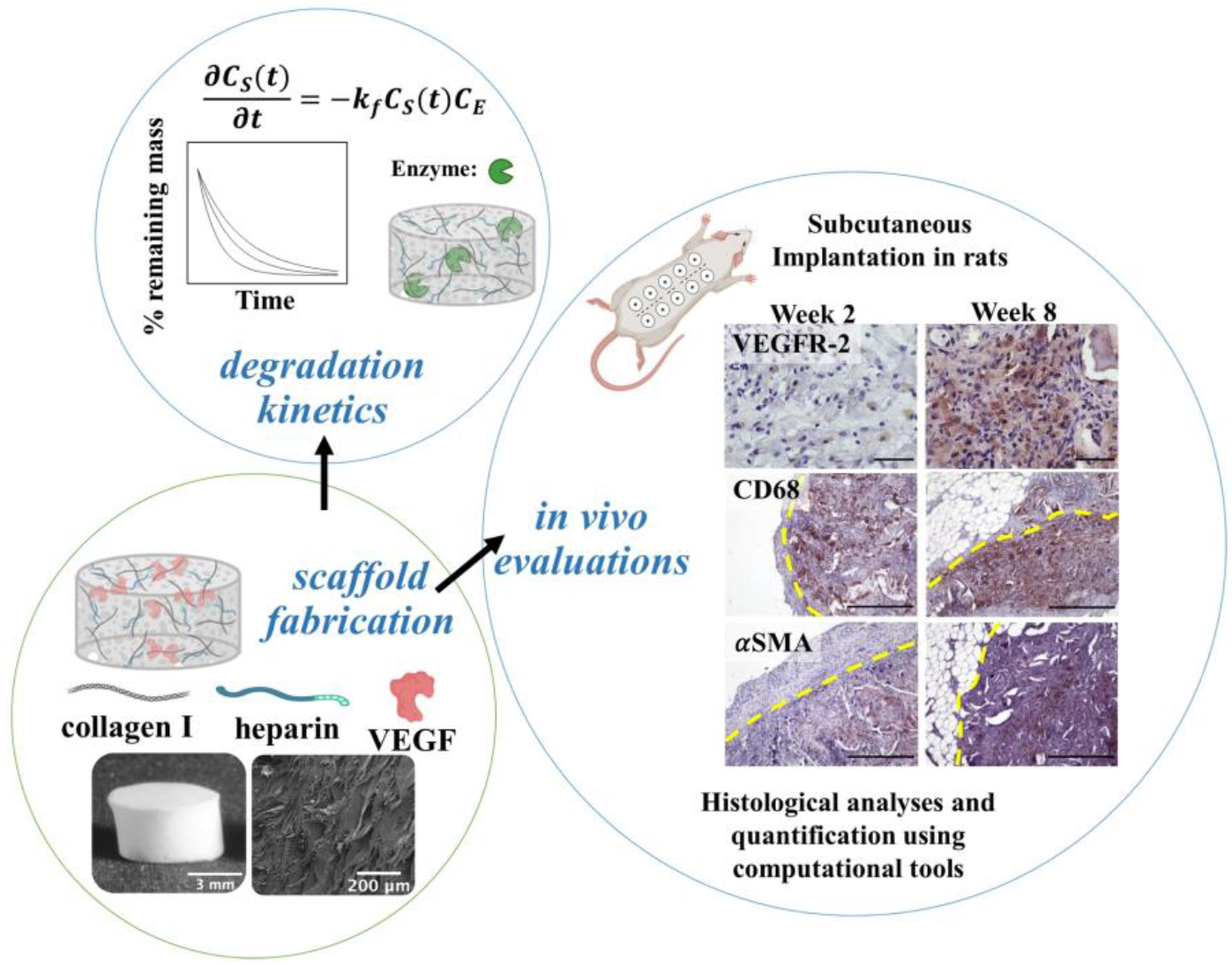

This work examines the role of scaffold fabrication and bioactive molecule inclusion on the enzymatic degradation of silk fibroin-based lyophilized sponges. Specifically, the roles of collagen I, heparin, and vascular endothelial growth factor are analyzed to determine the impact of formulation on rate of degradation. In addition, scaffolds are either pre-fabricated, where these bioactive molecules are included in the polymer solution prior to casting the scaffold or the bioactive molecules are introduced following scaffold formation through passive adsorption to the silk fibroin scaffold surface. Scaffolds are enzymatically degraded *in vitro*, and kinetic rate constants are calculated for the different formulations. *In vivo*, cellularity, adipose tissue accumulation, and scaffold area are assessed over time. Additionally, immunohistochemistry is used to visualize VEGF Receptor 2, CD 68, and α-smooth muscle actin over time.

## 1. Introduction

There is a need for biomaterials to supplement surgical procedures and aid in tissue and organ reconstruction. Procedures involving tumor resection, correction of anatomic congenital defects, and repair of traumatic or ischemic injuries benefit from *supportive* surgical materials. Defining what *supportive* means across patient populations with a variety of co-morbidities and complications is not trivial. Currently, the materials available to surgeons^7-8^ are not specific to patient characteristics (*e*.*g*., age, healing rate) and are not crafted to adapt to various injured environments or individual complications.

Investigating the rate of scaffold biodegradation and release of growth factors is essential to build adaptable materials for individualized needs.^9-10^ To facilitate neo-tissue formation and healing at the injury or surgical site, the surgical support material, or biomaterial, should degrade at a rate consistent with neo-tissue formation, while providing the required signaling modalities for healthy wound healing.^11-12^ However, the rate and quality of new tissue formation is highly dependent on the tissue location as well as patient-specific factors regarding their overall health, such as underlying co-morbidities, exposure to environmental toxins, or states of chronic inflammation.^13-16^ Accordingly, implantable biomaterials should involve tunable biodegradation and controllable bioactive molecule delivery systems to match patient-specific needs.

Currently, silk fibroin (silk) protein has been widely explored for *in vitro* tissue modeling,^17-18^ tissue replacement with clinical potential,^19^ and bioactive molecule or pharmaceutical delivery.^20-21^ Silk biomaterials are naturally-derived and degrade via proteolytic enzymes, resulting in small peptides or amino acids that can be absorbed by metabolically active stromal or immune cells. Previous work has examined the rate of degradation of a wide variety of silk-based materials *in vitro* using enzymes such as proteinase K, protease XIV, collagenase, and matrix metalloproteinase-1 (MMP-1).^3, 5, 22-26^ These results show that the molecular weight of silk, the concentration of silk in solution, the methods used to induce silk protein crystallinity, and the pore size of the material allow engineers to control or tune the rate of degradation. Ultimately, degradation studies of silk biomaterials can be utilized to match the timescale of the wound healing cascade for each patient and injury.^26^ It is hypothesized that *in vivo* degradation is a function of both enzyme-scaffold interaction and cellular infiltration. However, predictable silk degradation kinetics and determining what original biomaterial formulations drive *in vivo* degradation rates are unknown.

Silk biomaterials can be designed with an array of biological signaling modalities that can be harnessed in biomaterial design to facilitate regenerative outcomes. These biological signaling modalities include the ability to easily enhance the silk biomaterial with relevant ECM components.^27-30^ This is important as the composition of the ECM can modulate the time scales over which growth factors can bind and remain sequestered in the local environment.^31-33^ In silk-based biomaterials with a subcutaneous rodent implant model, aligned lyophilized silk sponges with the addition of decellularized cardiac ECM led to cell infiltration and promoted vascularization when compared to aligned lyophilized silk sponges alone.^28^ Using a silk hydrogel with the addition of decellularized cardiac ECM in a subcutaneous injection rodent model also led to cellular infiltration and promoted endothelial cell growth.^30^ The exact timing and release of key factors from the extracellular space to cells remains to be identified in order to optimize cellular responses.

In addition to ECM proteins, glycosaminoglycans (GAGs), such as heparin, aid in the local sequestration of growth factors within the complex ECM network and can modulate release of these factors as a function of tissue remodeling and repair.^34-35^ Consequently, heparin is often incorporated into tissue engineered constructs to mimic the extracellular environment.^36-39^ Heparin is an United States Food and Drug Administration approved medical product and can bind growth factors through specific electrostatic interactions to protect them from enzymatic degradation.^40-42^ Specifically, sulfate groups on heparin can bind to the positively charged regions on growth factors and other proteins. *In vitro, v*ascular endothelial growth factor (VEGF) and heparin binding interactions have been extensively explored.^43-44^ VEGF plays a key role in angiogenesis which is critical for tissue regeneration and repair following injury.^45-48^ By binding to VEGF receptors 1 and 2 (VEGFR-1 and VEGFR-2), VEGF stimulates endothelial cells. VEGF signaling through VEGFR-1 can also recruit monocytes and macrophages to the injured area,^49-50^ leading to a shift in matrix metalloproteinase (MMP) release. This results in ECM degradation and remodeling. On the other hand, prolonged VEGF expression can alter macrophage polarization and interleukin-10 (IL-10) secretion, provoking disruptive vessel growth.^51^ The ability of ECM and GAGs to sequester signals provides opportunities to adjust release rates of these signals.

Recent work has also shown that adipose tissue produces both pro- and anti-inflammatory cytokines and other biological mediators.^52-53^ Adipose tissue is an organ consisting mainly of adipocytes and is a triglyceride storage depot for regulation of energy homeostasis. During fasting and exercise, triglycerides are mobilized to release fatty acids into circulation in accordance with the body’s energy needs. Significant triglyceride accumulation in non-adipose tissues and organs, such as skeletal muscle and liver, can compromise their physical functions and insulin resistance. In addition to adipocytes, adipose tissue also contains preadipocytes, fibroblasts, vascular endothelial cells, and an assortment of immune cells. The infiltration of proinflammatory immune cells is associated with an increased production of proinflammatory cytokines including tumor necrosis factor α, IL-1β and IL-6.^54-57^ As a result, biomaterial researchers need to identify the extent of adipose tissue deposition within, and around degrading biomaterials for specific applications^57-60^ and the role this adipose tissue plays in final tissue or organ function.

Silk biomaterials have been fabricated with different functionalization techniques to control secreting factors.^61^ Two functionalization techniques include pre (-) fabrication loading and post (+, _S_) fabrication coatings.^62-64^ Pre (-) fabrication loading includes directly mixing the bioactive molecules in silk solution before biomaterial formation. Post (+, _S_) fabrication coatings are carried out by soaking the silk biomaterial in soluble solutions of bioactive molecules. Using pre (-) fabrication loading techniques, silk biomaterials can maintain bioactivity of the stabilized bioactive molecules. A drawback of this technique is it results in loading inefficiencies or loss of the original mass incorporated. This is a function of the numerous processing steps to create some silk biomaterials or sterilization techniques for *in vivo* translation. Post (+, _S_) fabrication coating can resolve these issues by soaking pre-processed silk biomaterials in soluble solutions of bioactive molecules. Our lab specifically focuses on the use of silk sponge-like scaffolds that are generated from freezing, lyophilizing, and autoclaving or water annealing *Bombyx mori* silk fibroin solution. It is hypothesized that using the pre (-) fabrication loading method will enable entrapment of bioactive molecules within the β-sheets of the lyophilized sponge-like scaffolds. Conversely, the post (+, _S_) fabrication method will allow bioactive molecule solutions to interact with the hydrophobic and hydrophilic parts of silk fibroin and not be trapped within the β-sheets of the lyophilized scaffolds.

This work focuses on the utilization of acellular biomaterials to instruct local cell behavior by controlling the scaffold fabrication method and composition. Here, we have developed an acellular sponge-like scaffold containing silk, type I collagen (collagen I), heparin, and VEGF. The simplicity of the system supports biological predictability to improve our understanding of protein-protein interactions related to regenerative medicine. The silk system with directing factors were fabricated via two methods to quantify the specific interactions between scaffold components. In subcutaneous implants into Sprague Dawley rats, the degradation of the scaffolds was dictated by the fabrication method and scaffold formulation. The percentage of cells infiltrating the scaffold increased when type I collagen, heparin, and/or VEGF were added to the scaffold. To gain further insight into the influence of each component and fabrication method, *in vitro* degradation rates of these scaffolds were assessed using proteinase K and protease XIV. Scaffold degradation occurred more rapidly with higher enzyme concentrations and increased theoretical cleavage sites. However, the addition of type I collagen and heparin did not change degradation rates when compared to the silk scaffold, suggesting that *in vitro*, addition of low concentrations of additional proteins doesn’t affect degradation rates. These *in vitro* results were contrary to observations *in vivo*, where ECM additions influence local ECM deposition, rates of immune cell infiltration, and vascular structure formation. Increases in adipose tissue in and around the degrading scaffold were observed over 8 weeks.

## 2. Results & Discussion

To promote desirable and functional recovery, a biomaterial needs to balance the regeneration of tissue with scaffold degradation, as well as mimic the native extracellular environment and act as a reservoir for growth factor binding and sequestration. Current methodologies for determining *in vivo* biomaterial performance include iterative biomaterial design through *in vitro* studies and characterization and subsequent *in vivo* surgical procedures. We aim to reduce the need for trial-and-error biomaterial design through an in-depth analysis of the parameters and fabrication strategies that drive *in vivo* performance, supported by mechanistic *in vitro* studies. We explored *in vitro* and *in vivo* analyses to determine drivers of *in vivo* metrics: scaffold degradation and cell infiltration. *In vivo* studies provide a holistic view of scaffold degradation, immune cell infiltration, adipose tissue deposition, and angiogenesis. *In vitro* studies were used to determine the mechanism of scaffold degradation based on protein-protein interactions in a controlled environment.

### 2.1 Scaffold fabrication and composition impacts *in vivo* performance over 8 weeks

The degradation of silk biomaterials *in vivo* relates to the protease-secreting cells often associated with host immune response. Immune cells, such as macrophages, can secrete matrix metalloproteinases (MMPs) that degrade components of the ECM and the silk biomaterial.^65^ We investigated how bioactive molecule incorporation altered *in vivo* performance in a subcutaneous rodent model using two fabrication methods, termedpre (-) fabrication (denoted by -) and post (+, _S_) fabrication (denoted by + and _S_) (**Figure 1**). The formulations and their use within this manuscript are organized in **Table 1**. Individual scaffold names, abbreviations, and associated symbols are shown in **Table S1**. To determine the effect of an individual bioactive molecule and/or fabrication method, we split the 10 conditions into groups according to their properties for data analysis (**Table 1**). Since it is hypothesized that *in vivo* degradation is a function of combined feedback between enzyme-scaffold interaction and cellular infiltration, we quantified the scaffold area and cellular infiltration using the description of variables from **Figure 2** and mathematical expressions detailed in **equations (1)** and **(2)**, respectively. These graphs are presented in **Figure 3** and discussed in detail below. Representative hematoxylin and eosin (H&E) and Masson’s Trichrome images used for the calculations for all conditions and time points are given in **Figures S1-S6**. Pre (-) fabrication, post (+, _S_) fabrication, and both fabrication method H&E images are shown in **Figure S1, Figure S2**, and **Figure S3**, respectively. Pre (-) fabrication, post (+, _S_) fabrication, and both fabrication method Masson’s Trichrome images are shown in **Figure S4, Figure S5**, and **Figure S6**, respectively.

**Figure 1.**
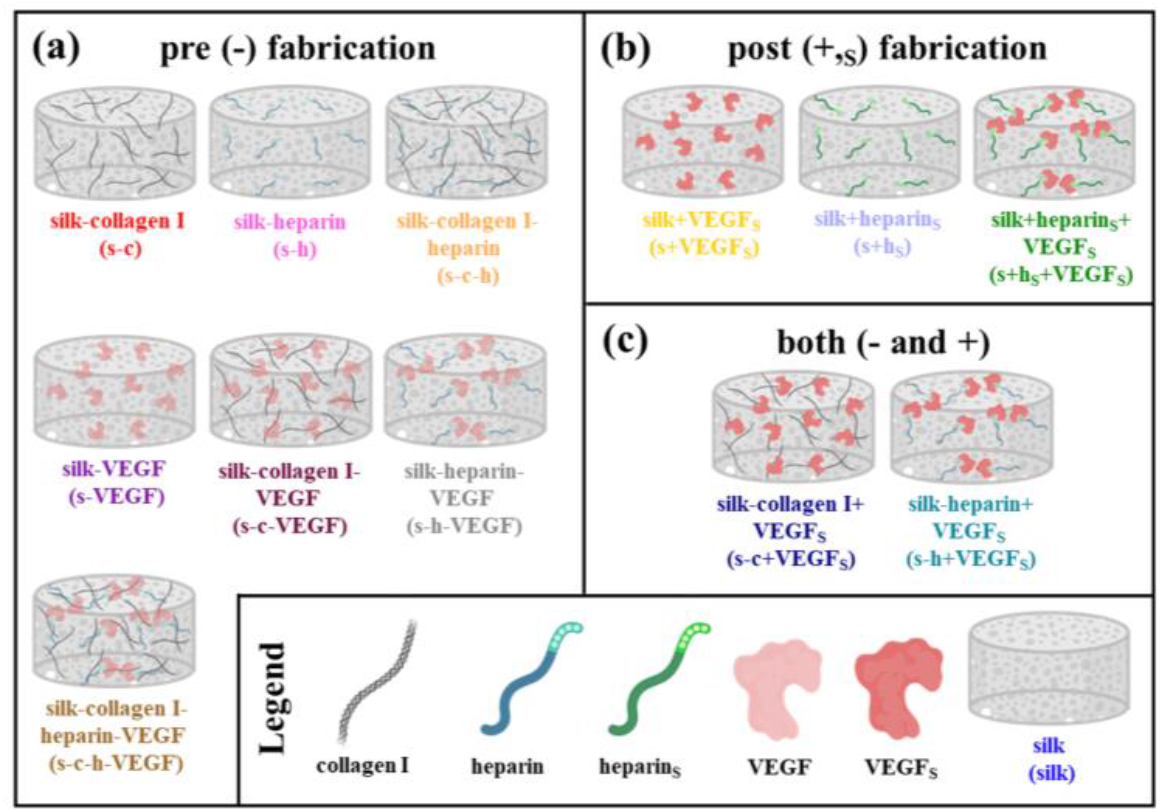
Schematic representation of scaffold formulations used for scaffold evaluation. (a) Pre (-) scaffolds were fabricated by including heparin (100 mg/mL), type I collagen (200 mg/mL), and/or VEGF (0.1 mg/mL) into the diluted aqueous silk solution. Following incorporation, the silk solution was utilized to create lyophilized scaffolds. (b) Post (+,_S_) fabrication scaffolds were made by soaking silk lyophilized scaffolds in solubilized solutions of heparin and VEGF for 30 min. VEGF_S_ and heparin_S_ designates soluble VEGF and heparin solutions added post (+, _S_) fabrication, respectively. (c) Two scaffolds were made using both the pre (-) fabrication method and the post (+,_S_) fabrication method, this is termed both (-,+) fabrication methods. For *in vivo* experiments, scaffolds had a radius of 3 mm and height of 3 mm. For *in vitro* experiments, scaffolds were cut to a radius of 3 mm and height of 4 mm.

**Table 1:** Scaffold terminology and scaffold groups for *in vivo* investigations and analysis. Columns identify data splits for statistical analysis presented in subsequent figures. With or containing the bioactive molecule is denoted by “with” (w/) while samples that do not contain that bioactive molecule are denoted as “without” (w/o).

| Scaffold formulation | pre (-) fabrication | post (+,s) fabrication | both fabrications (-,+) | w/ collagen I | w/o collagen I | w/ heparin | w/o heparin | w/ VEGF | w/o VEGF |
| --- | --- | --- | --- | --- | --- | --- | --- | --- | --- |
| **silk** |  |  |  |  |  |  |  |  |  |
| **silk-collagen I** | √ |  |  | √ |  |  | √ |  | √ |
| **silk-heparin** | √ |  |  |  | √ | √ |  |  | √ |
| **silk-heparin-VEGF** | √ |  |  |  | √ | √ |  | √ |  |
| **silk-collagen I-heparin-VEGF** | √ |  |  | √ |  | √ |  | √ |  |
| **silk+VEGF<sub>s</sub>** |  | √ |  |  | √ |  | √ | √ |  |
| **silk+heparin<sub>s</sub>** |  | √ |  |  | √ | √ |  |  | √ |
| **silk+heparin<sub>s</sub>+VEGF<sub>s</sub>** |  | √ |  |  | √ | √ |  | √ |  |
| **silk-collagen I+VEGF<sub>s</sub>** |  |  | √ | √ |  |  | √ | √ |  |
| **silk-heparin+VEGF<sub>s</sub>** |  |  | √ |  | √ | √ |  | √ |  |

**Figure 2.**
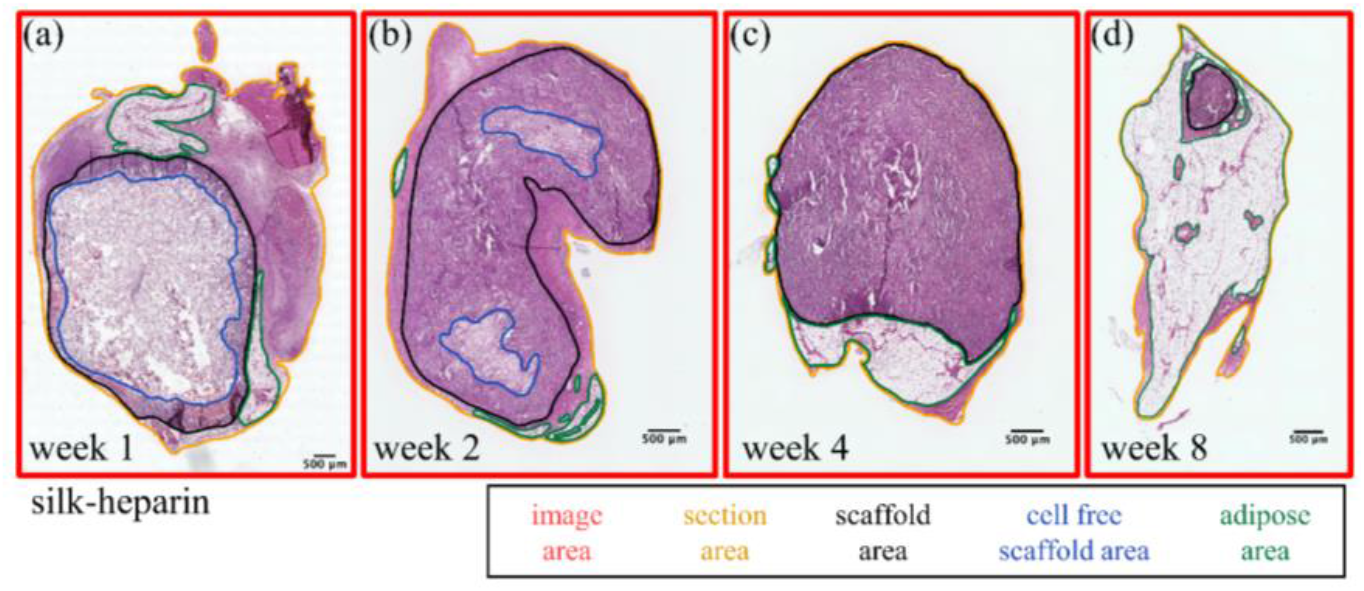
Characterization of *in vivo* scaffold dimensions with H&E images for silk-heparin biomaterials at (a) week 1, (b) week 2, (c) week 4, and (d) week 8. The red outline is the image area. The orange outline is the section area. The black outline is the scaffold area. The blue outline is the cell free area of the scaffold. The green outline is the adipose area.

**Figure 3.**
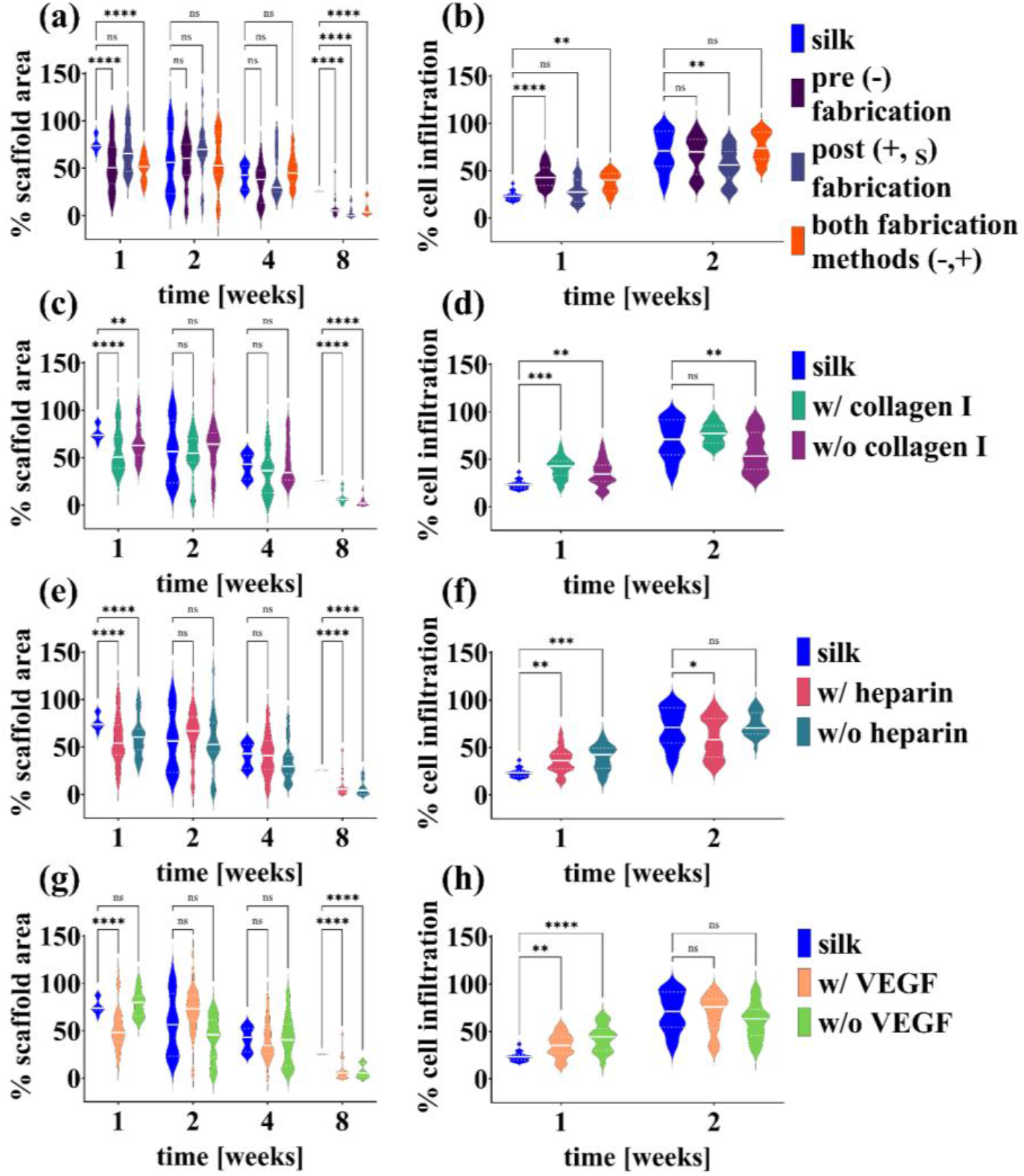
Quantitative changes of scaffold area (a,c,e,g) and cell infiltration (b,d,f,h) with scaffolds made from different fabrication methods (a, b) and with different bioactive molecules (c-h) after explant from a subcutaneous rodent model are presented here. Scaffolds were explanted 1-, 2-, 4-, and 8-weeks post-surgery. Scaffold area % and cell infiltration % were calculated with the description of variables in Figure 2 and mathematical expressions detailed in equation (3) and (4), respectively. Statistical significance was determined using a mixed-effects model with Dunnett’s post-hoc multiple comparisons test. Statistical significance is reported as *p < 0.05, **p < 0.01, ***p < 0.001, and ****p < 0.0001.

#### 2.1.1 Scaffolds formed by pre (-) fabrication methods or both fabrication methods (+,-) decrease scaffold area at early time points

To explore the role of scaffold fabrication on *in vivo* scaffold degradation and cell infiltration, we split the data into 3 groups: pre (-) fabrication, post (+,_S_) fabrication, or both fabrication methods (+,-). We then analyzed H&E and Masson’s Trichrome images of explanted scaffolds to determine scaffold area and cell infiltration over time. The description of variables in **Figure 2** and the mathematical expressions detailed in **equations (1)** and **(2)** were used to calculate the % scaffold area and % cell infiltration. We found that using the pre (-) fabrication method or both fabrication methods (+,-) for scaffold formation decreased the scaffold area at early time points (week 1) (**Figure 3a**). At weeks 2 and 4, there was no significant difference in the scaffold area regardless of scaffold fabrication method as compared to neat silk scaffolds (**Figure 3a**). By week 8, we find that using any fabrication method significantly decreases the scaffold area compared to neat silk scaffolds (**Figure 3a**).

#### 2.1.2 Pre (-) fabrication formed scaffolds have rapid and persistent cell infiltration

At week 1, cell infiltration was significantly higher for scaffolds made from any fabrication method as compared to neat silk scaffolds (**Figure 3b**). It is important to note that the scaffolds formed by the pre (-) fabrication method had the highest % cell infiltration (**Figure 3b**). By week 2, the post (+,_S_) fabrication method scaffolds had a significantly lower % cell infiltration than did the neat silk scaffolds (**Figure 3b**). By week 8, all scaffolds were completely cellularized and thus no differences were noted.

In this study design, we hypothesized that the pre (-) fabrication method would be advantageous due to the interactions of the silk protein with the bioactive components prior to the formation of β-sheet structures, allowing the bioactive molecule to be entrapped within the silk protein structure. The hypothesized advantage of the pre (-) fabrication method is that it can lead to growth factor stability within the lyophilized sponge-like scaffold, prolonging bioactivity. The post (+,_S_) fabrication method hypothesizes that the soluble heparin and VEGF proteins will interact with the hydrophobic and hydrophilic parts of silk fibroin fibrous structure, formed after crystallization has taken place. Thus, the proteins imbibed into the scaffolds using the post (+,_S_) fabrication method will not be trapped within the β-sheets of the lyophilized scaffolds. We hypothesized that these two methods may alter the interactions of the biomaterial with the host tissue. Studies employing the pre (-) fabrication method have shown improved cell infiltration when ECM components are incorporated into silk scaffolds.^28^ Studies employing the post (+,_S_) fabrication method have also seen increased cell infiltration.^66^ However, *in vitro* studies observed that increasing the soaking time can increase cell attachment where the highest cell attachment occurred with 7 days of soaking the silk scaffold.^64^

Our results demonstrate that at week 1, the post (+,_S_) fabrication method does increase cell infiltration compared to neat silk scaffolds (**Figure 3b**), which is in agreement with previous studies.^66^ By week 2, the scaffolds formed with the post (+,_S_) fabrication method have the lowest cell infiltration. This indicates that the heparin and/or VEGF have been degraded or have fully diffused out of the scaffold, and concentrations remaining within the scaffold are not sufficient to elicit a change in cell response. Broader comparisons with different scaffold soaking times^64^ with soluble heparin and VEGF may impact the results observed here. Despite potential impact for improvement with longer soaking times, the pre (-) fabrication method eliminates the need for this additional step as demonstrated by high levels of cell infiltration and decreased scaffold area over time (**Figure 3a,b**).

#### 2.1.3 Addition of type I collagen, heparin, and/or VEGF leads to decreased scaffold area

To investigate how the addition of bioactive molecules impacts *in vivo* scaffold degradation and cell infiltration, we analyzed H&E and Masson’s Trichrome images of explanted scaffolds made with collagen I, heparin, and/or VEGF. The description of variables in **Figure 2** and the mathematical expressions detailed in **Equation (1)** and **Equation (2)** were used to calculate the % scaffold area and % cell infiltration. **Figure 3c** shows the scaffold area over 8 weeks for neat silk scaffolds compared to the full data set split into two categories: scaffolds with collagen I and scaffolds without collagen I (**Table 1**). At week 1, neat silk scaffolds had significantly higher scaffold areas when compared to either scaffolds with collagen I or scaffolds without collagen I. By weeks 2 and 4, there was no significant difference between silk scaffolds and scaffolds with or without collagen I. By week 8, silk scaffold areas became significantly greater than scaffolds with or without collagen I, again following the same trend as week 1, where addition of any bioactive molecule led to increased rates of scaffold degradation.

Again, we perform a data split of the full data set to compare scaffolds containing heparin to those without heparin, regardless of the presence of other bioactive molecules or the fabrication method (**Table 1**). **Figure 3e** shows the scaffold area over 8 weeks for silk scaffolds, scaffolds with heparin, and scaffolds without heparin. At weeks 1 and 8, neat silk scaffolds had significantly higher scaffold areas (less degradation) when compared to either scaffolds with heparin or scaffolds without heparin. At weeks 2 and 4, there was no significant difference between neat silk scaffolds and scaffolds with or without heparin.

Similarly, another data split was performed to assess the role of VEGF. **Figure 3g** shows the scaffold area over 8 weeks for silk scaffolds, scaffolds with VEGF, and scaffolds without VEGF. At week 1, silk scaffolds with VEGF had significantly smaller scaffold areas than neat silk scaffolds, showing greater degradation at early timepoints with VEGF addition. At weeks 2 and 4, there was no significant difference between neat silk scaffolds and scaffolds with or without VEGF. By week 8, neat silk scaffolds again had significantly higher scaffold areas when compared to either scaffolds with VEGF or scaffolds without VEGF. The experiments presented here indicate that the addition of a biological molecule increases scaffold degradation in a subcutaneous implant model. When looking at positive and negative controls independently (**Figure S7, S8, S9**), we further confirm that the addition of collagen I, heparin, or VEGF can decrease scaffold area. Previous researchers have also observed that the addition of bioactive molecules increases scaffold degradation *in vivo*.^67-69^ Schirmer *et al*. have subcutaneously implanted GAG-based hydrogels with either VEGF, fibroblast growth factor 2 (FGF2, bFGF), or stromal cell-derived factor (SDF1α) and, after 28 days, hydrogels with any growth factor were significantly more degraded than unloaded controls,^68^ similar to our results. However, they did not show any difference in degradation after 1 week. Together, data presented in **Figures 3c,e,g** show that the addition of any bioactive molecule increases scaffold degradation as noted by the increased scaffold area for neat silk scaffolds comparatively, regardless of how the scaffold was formed or what it contained. This suggests that the addition of any bioactive molecule impacts scaffold degradation, necessitating investigation of cell infiltration and the types of cells present within the scaffold over time.

#### 2.1.4 Rapid *in vivo* cell infiltration occurs when scaffolds are formed with the addition of a biological molecule

The same data splits are used to analyze cell infiltration, but by week 4, all scaffold formulations were fully infiltrated (>95% cell infiltration calculated from data in **Figures S10-12**). **Figure 3d** shows the % cell infiltration for neat silk scaffolds, scaffolds with collagen I, and scaffolds without collagen I. At week 1, neat silk scaffolds had significantly lower % cell infiltration than did scaffolds with collagen I or scaffolds without collagen I. At week 2, there was no significant difference in % cell infiltration between neat silk scaffolds and scaffolds with collagen I. However, at week 2, scaffolds without collagen I had significantly lower % cell infiltration compared to neat silk scaffolds. By week 4, all scaffold formulations had >95% cell infiltration. **Figure 3f** shows the % cell infiltration for silk scaffolds, scaffolds with heparin, and scaffolds without heparin. At week 1, neat silk scaffolds had significantly lower % cell infiltration than did scaffolds with heparin or scaffolds without heparin. At week 2, there was no difference in % cell infiltration between groups. By week 4, all scaffolds had >95% cell infiltration. **Figure 3h** shows the % cell infiltration for silk scaffolds, scaffolds with VEGF, and scaffolds without VEGF. At week 1, silk scaffolds had significantly lower % cell infiltration than did scaffolds with VEGF or scaffolds without VEGF. At week 2, there was no significant difference in % cell infiltration between groups. Again, by week 4, all scaffolds had >95% cell infiltration regardless of the bioactive molecule. These results suggest that collagen I, heparin, or VEGF, when analyzed by this data split strategy, are not singly the major driver variation in cell infiltration.

Experimentally, we were able to determine that the addition of any biological molecule (collagen I, heparin, VEGF) increased cell infiltration at early time points. This was determined as at week 1 all groups (scaffolds w/collagen I, w/o collagen I, w/heparin, w/o heparin, w/VEGF, w/o VEGF) had significantly higher cell infiltration than did neat silk scaffolds. Since groups w/o collagen I, w/o heparin, and w/o VEGF still had other biological molecules within the scaffold, the conclusion can be made that the addition of any bioactive molecule increased cell infiltration. Additionally, when looking at positive and negative controls independently (**Figure S10, S11, S12**), we further confirm that the addition of collagen I, heparin, or VEGF can positively influence cell infiltration.

Previous studies have shown that the addition of ECM components can increase cell infiltration in various injury environments.^7, 70-72^ As mentioned previously, the addition of decellularized cardiac ECM to align lyophilized silk scaffold or silk hydrogels resulted in increased cellular infiltration and promoted vascularization in comparison with neat silk samples.^28, 30^ Another study found that heparin-based coacervate systems that were enhanced with FGF2 displayed significantly accelerated wound healing and increased cell proliferation compared to the control coacervate alone.^70^ Researchers also found that the introduction of a vascular-inducing peptide into a silk fibroin hydrogel increased cellular infiltration and doubled vascularization in comparison with the hydrogel alone.^72^ Our results stating that the inclusion of biological molecules into the scaffold increases cellular infiltration are in agreement with the current literature, but were surprising as we anticipated being able to more granularly identify drivers of scaffold infiltration with this experimental design.

Assessing and quantifying scaffold degradation and cellular infiltration *in vivo* is not standardized within the field. Hence, there are limitations to our current analyses that may yield greater error in the results and compromise our ability to detect slight differences between formulations, as we had anticipated. For example, slight variations in scaffold preparation for histology may increase the error in calculations such as scaffold area or % cell infiltration, given that these are area-based measurements. More specifically, sectioning challenges can arise and shifts in the section on the glass slide during staining and mounting could alter the size of the section leading higher error in the data.

The post (+, _S_) fabrication method poses challenges for *in vitro* analysis due to the labile nature of the bioactive molecules within the scaffold and the role diffusion of these molecules out of the scaffold may play in observed *in vivo* results. Thus, for the remainder of the paper we will focus on pre (-) fabrication methods for incorporating bioactive molecules into lyophilized scaffolds both due to the increased potential for translation of these formulations and the observed improvements in initial cell-scaffold interactions.

### 2.2 Using *in vitro* data to model scaffold degradation

#### 2.2.1 Addition of collagen I and heparin to scaffolds does not change *in vitro* degradation

After seeing disparities in scaffold area and cell infiltration *in vivo*, we aimed to further probe the driving force for scaffold degradation. Two major hypotheses exist: 1) the scaffold formulation dictates the rate of degradation based on the available cleavage sites or 2) the scaffold formulation dictates the recruitment of infiltrating cells which in turn lead to observed spatiotemporal changes in scaffold area and rate of cell infiltration. To investigate the first hypothesis, we sought to model the degradation of the acellular scaffolds *in vitro* in a cell-free environment. We compared the use of modified first-order reaction kinetics vs Michaelis-Menten reaction kinetics^26^ in describing experimental results of isotropic, lyophilized, composite silk scaffold degradation. Consistent with previously reported results,^26^ a modified first-order kinetic model was appropriate for kinetic rate constant estimation from experimental degradation data (**Figure 4**), even when accounting for the errors in the experimental data. **Table 2** shows the results for modified first-order rate constant estimation and the accompanying standard deviation from Monte Carlo studies for silk, s-c, s-h, and s-c-h scaffolds degraded using proteinase K (1 U/mL and 0.1 U/mL).

**Figure 4.**
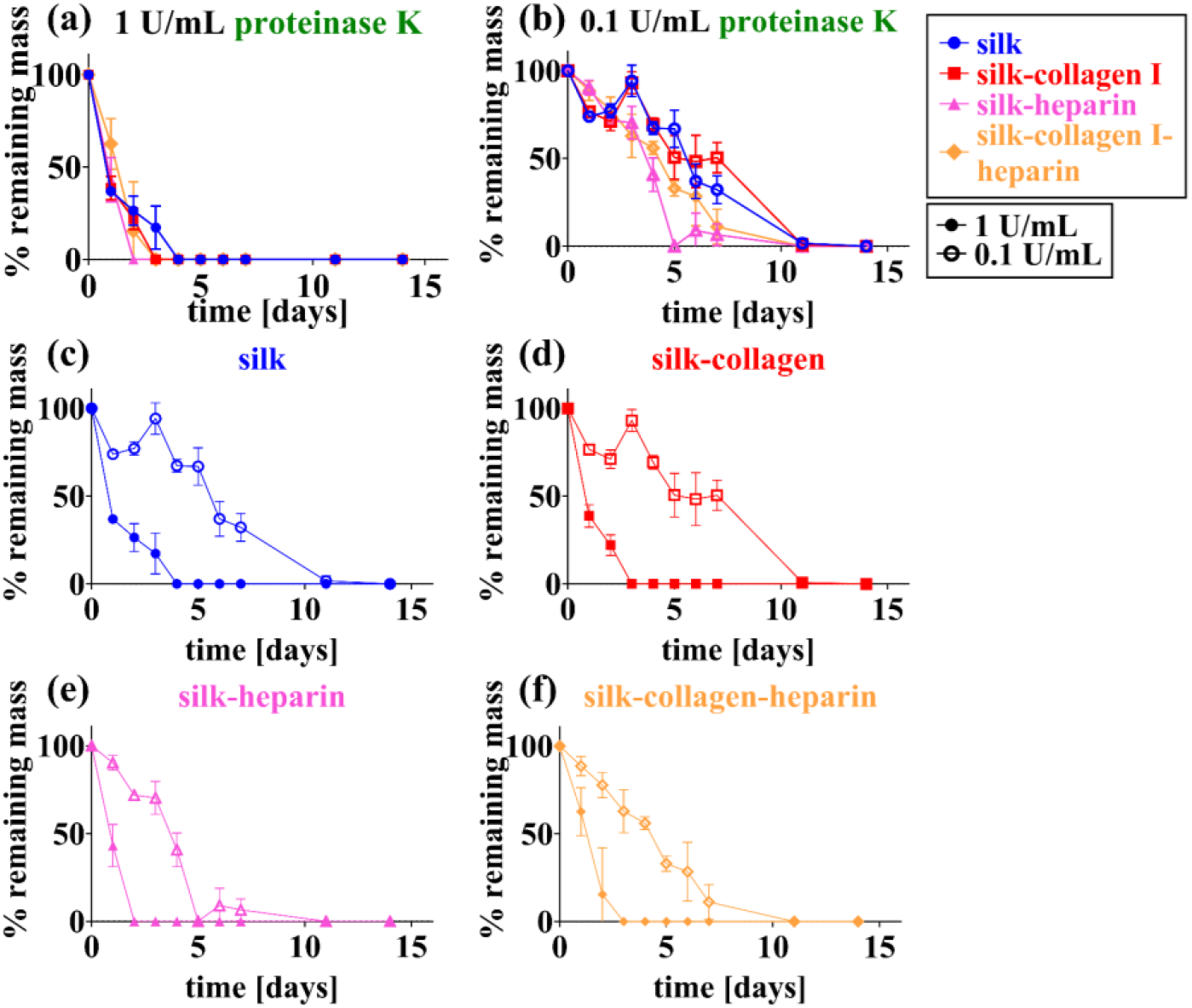
Degradation of biomaterials is necessary for *in vivo* tissue regeneration. Quantitative changes of silk scaffold mass with different bioactive molecules during enzymatic degradation are illustrated here. Silk, silk-collagen I, silk-heparin, and silk-collagen I-heparin were immersed in (a) proteinase K (1 U/mL) and (b) proteinase K (0.1 U/mL) and incubated at 37°C. The addition of collagen I and/or heparin did not change the rate of degradation of the composite silk scaffolds over 28 days for proteinase K for all concentrations (1 U/mL, 0.1 U/mL). A higher concentration of proteinase K (1 U/mL) degraded silk (c), s-c (d), silk-heparin (e), and silk-collagen I-heparin (f) scaffolds faster than 0.1 U/mL proteinase K. The ordinate on all graphs is expressed as the mass of the sample divided by the starting point mass multiplied by 100. Data expressed by mean ± 1 SD with 3 biological replicates.

**Table 2:** Modified first order rate constant (*k*_*f*_) for 12 hr WA silk composite scaffolds immersed in varying concentrations of proteinase K. Where *k*_*f*_ is determined from weighted least squares analysis and standard deviation for *k*_*f*_ comes from Monte Carlo studies.

| scaffold | enzyme | enzyme concentration [U/mL] | $k_f$ [mL/U·day] |
| --- | --- | --- | --- |
| silk<sup>a</sup> | proteinase K | 1 | 1.00 ± 0.11 |
| s-c | proteinase K | 1 | 1.06 ± 0.13 |
| s-h | proteinase K | 1 | 0.98 ± 0.51 |
| s-c-h | proteinase K | 1 | 0.59 ± 0.24 |
| silk<sup>a</sup> | proteinase K | 0.1 | 2.20 ± 0.75 |
| s-c | proteinase K | 0.1 | 1.75 ± 0.59 |
| s-h | proteinase K | 0.1 | 1.65 ± 0.27 |
| s-c-h | proteinase K | 0.1 | 1.34 ± 0.39 |

<sup>a</sup>Data obtained from Jameson et al. for comparison.

Silk-based scaffolds with collagen I and heparin were explored to elucidate the role of the addition of other biopolymers on silk scaffold degradation with proteinase K (1 U/mL and 0.1 U/mL) (**Figure 4a-b**). When immersing scaffolds in 1 U/mL proteinase K, the % remaining mass of silk scaffolds became statistically similar to 0% mass or 0 mg at day 3, the % remaining mass of s-c scaffolds became statistically similar to 0% mass or 0 mg at day 3, the % remaining mass of s-h scaffolds became statistically similar to 0% mass or 0 mg at day 2, and the % remaining mass of s-c-h scaffolds became statistically similar to 0% mass or 0 mg at day 2 (**Figure 4a**). Experimentally, we were able to determine the addition of bioactive molecules (collagen I and heparin) at low concentrations did not influence the rate of degradation for scaffolds immersed in 1 U/mL proteinase K. Additionally, the rate constants for proteinase K at 1 U/mL were similar across silk compositions with silk, s-c, s-h, and s-c-h having rate constants of *k*_*f*_ = 1.0 ± 0.11 mL/U·day, 1.06 ± 0.13 mL/U·day, 0.98 ± 0.30 mL/U·day, and 0.59 ± 0.24 mL/U·day, respectively (**Table 2**).

Lastly, scaffolds with small concentrations of bioactive molecules immersed in proteinase K at 0.1 U/mL also confirmed that the bioactive molecules did not change the rate of degradation. The % remaining mass from silk sponges became statistically similar to 0% remaining mass or 0 mg at day 11, silk-collagen I sponges became statistically similar to 0% remaining mass or 0 mg at day 11, silk-heparin sponges became statistically similar to 0% remaining mass or 0 mg at day 7, and silk-collagen I-heparin sponges became statistically similar to 0% remaining mass or 0 mg at day 7 (**Figure 4b**). Moreover, the rate constants for proteinase K at 0.1 U/mL were similar across silk compositions with silk, s-c, s-h, and s-c-h having rate constants of *k*_*f*_ = 2.2 ± 0.75 mL/U·day, 1.75 ± 0.59 mL/U·day, 1.65 ± 0.27 mL/U·day, and 1.34 ± 0.39 mL/U·day, respectively (**Table 2**).

In this *in vitro* study, we used proteinase K, which is a serine protease. These proteases cleave peptide bonds adjacent to aromatic, aliphatic, and hydrophobic amino acids. Proteinase K from *Engyodontim album* cleaves amide bonds adjacent to histidine, phenylalanine, tryptophan, tyrosine, alanine, isoleucine, leucine, proline, valine, and methionine (**Table 2**).^73-74^

The silk protein from *Bombyx mori* silk cocoons consists of a heavy chain, a light chain, and a P25 glycoprotein.^75-76^ The light chain and P25 glycoprotein account for only 5% of the amino acids present in silk.^75-76^ The heavy chain of silk is the major constituent and contains 11 small hydrophilic domains and 12 hydrophobic domains. The hydrophobic domains contain β-sheet forming regions with repetitive amino acid motifs consisting of glycine-X (GX) repeats where X can be alanine, serine, or tyrosine. The number of cleavage sites was predicted within the heavy and light chain of silk for proteinase K.^3-4^ The predicted values for proteinase K is ∼2200 (**Table 2**). This value does not consider the accessibility of the cleavage sites due to secondary structure.

Fibrous type I collagen contains an abundance of three amino acids—glycine, proline, and hydroxyproline—and forms a triple helical structure.^77^ This creates a repeating motif of glycine-proline-X, where X can be any amino acid.^77^ Upon investigating the number of histidine, phenylalanine, tryptophan, tyrosine, alanine, isoleucine, leucine, proline, valine, and methionine residues within collagen I, it is estimated there is ∼410 cleavage sites for proteinase K to degrade collagen I (Accession number: P02454), a number much lower than the ∼2200 cleavage sites predicted for silk fibroin (**Table 3**). Additionally, the mass ratio of silk to collagen I within our scaffolds is 150:1. Hence, the collagen I within our silk scaffolds is contributing no effect on the degradation rate of the scaffold when immersed in proteinase K.

**Table 3:** Proteolytic enzyme cleavage sites for the heavy and light chain of silk fibroin, collagen I, laminin, fibrinogen, and collagen IV. Data compiled from published resources.^1-6^.

| enzyme | cleavage sites | # of estimated cleavage sites |  |  |  |  | ratios |  |  |  |
| --- | --- | --- | --- | --- | --- | --- | --- | --- | --- | --- |
|  |  | silk fibroin (A) | collagen I (B) | laminin (C) | fibrinogen (D) | collagen IV (E) | (A):(B) | (A):(C) | (A):(D) | (A):(E) |
| proteinase K | His, Phe, Trp, Tyr, Ala, Ile, Leu, Pro, Val, Met | ~2200 | ~410 | ~470 | ~240 | ~625 | 5:1 | 5:1 | 9:1 | 4:1 |
| protease XIV | Tyr, Phe, Trp, His, Lys, Arg | ~390 | ~110 | ~300 | ~185 | ~255 | 4:1 | 1:1 | 2:1 | 2:1 |
| MMP-1 | Gly-Ile, Gly-Leu | ~10 | ~20 | ~5 | ~5 | ~65 | 1:2 | 2:1 | 2:1 | 1:7 |
| MMP-2 | Gly-Ile, Gly-Leu, Gly-Val, Gly-Phe, Gly-Asn, Gly-Ser | ~600 | ~75 | ~30 | ~50 | ~135 | 8:1 | 20:1 | 12:1 | 4:1 |

Heparin, an acidic polysaccharide in the glycosaminoglycan family, consists of a repeating disaccharide structure of 1→4-linked pyranosyluronic acid and glucosamine saccharide residues.^78-79^ Heparin is degraded by two enzymes, the prokaryotic polysaccharide lyases and the eukaryotic glucuronyl hydrolases.^80-81^ Prokaryotic polysaccharide lyases act through an eliminative mechanism whereas eukaryotic glucuronyl hydrolases act through a hydrolytic method.^80-81^ Although serine proteases catalyze the hydrolysis of peptide bonds, they do not catalyze the hydrolysis of glycosidic bonds found in polysaccharides. This can explain why there is no difference in the rate of degradation when heparin is added to the silk scaffolds.

#### 2.2.2 Enzyme concentration impacts scaffold degradation *in vitro*

Varying the concentration of enzyme solution was probed to determine the influence of enzyme concentration on scaffold degradation rate. We explored enzymatic degradation of scaffolds (silk, s-c, silk-heparin, and s-c-h) immersed in both 0.1 U/mL and 1 U/mL proteinase K (**Figure 4c-f**). As seen in **Figure 4c-f** and *k*_*f*_ values in **Table 2**, decreases in enzyme concentration increased the timeline for degradation.

The experimental data and rate constant estimation are as expected from previous research for all scaffold compositions (silk, silk-collagen I, silk-heparin, and silk-collagen I-heparin).^22, 26^ The addition of collagen I and/or heparin at low concentrations in the silk scaffold did not alter enzyme degradation. This data further confirms that there is silk scaffold substrate available to bind the enzyme increasing the rate of degradation. Nonetheless, at increasing enzyme concentrations there is a maximum at which the silk scaffold substrate will be bound fully, and the rate of degradation will not alter with a higher enzyme concentration. These high enzyme concentrations are not expected *in vivo*.

Using the analysis from *in vitro* experiments of enzymatic degradation provides a new understanding of the *in vivo* work presented. From the *in vitro* experiments, the enzyme concentration is the major parameter dictating scaffold degradation. The addition of collagen I and heparin to the scaffold did not change the rate of degradation *in vitro*. However, when using the scaffold area as a measurement of *in vivo* degradation, there are some differences in scaffold area when biological components (collagen I, heparin, and/or VEGF) were included into the scaffold (**Figure 3a-d**). As expected, these results suggest that the mechanism behind *in vivo* degradation is different than *in vitro*. The addition of collagen I, heparin, and/or VEGF could differentially recruit cell types or amounts. Differences in cell types can lead to shifts in enzyme secretion, specifically MMPs. MMPs are a major class of zinc-containing and calcium-dependent endoproteinases and are instrumental for physiological processes such as tissue remodeling, wound healing, angiogenesis, and organogenesis.^82-83^ MMPs degrade ECM proteins and are secreted by a variety of cell types including monocytes and macrophages.^84^ Two common MMPs, MMP-1 and MMP-2, cleave specific peptides (MMP-1: Gly-Ile, Gly-Leu, MMP-2: Gly-Ile, Gly-Leu, Gly-Val, Gly-Phe, Gly-Asn, Gly-Ser).^1-2^ Brown *et al*. predicted the number of cleavage sites within the heavy and light chain of silk fibroin for MMP-1 and MMP-2 to be ∼10 and ∼600, respectively (**Table 3**).^3^ It is estimated that there is ∼20 cleavage sites for MMP-1 to degrade collagen I, and ∼75 cleavage sites for MMP-2 to degrade collagen I (Accession number: P02454, **Table 3**). To expand our results to different ECM components, we predicted the number of cleavage sites for laminin (Accession number: P11047), fibrinogen (Accession number: P02671, and collagen IV (Accession number: P53420, **Table 3**). It is estimated that there is ∼5 cleavage sites for MMP-1 to degrade both laminin and fibrinogen while there is ∼65 cleavage sites for MMP-1 to degrade collagen IV. Additionally, it is estimated there is ∼30 cleavage sites for MMP-2 to degrade laminin, ∼50 cleavage sites for MMP-2 to degrade fibrinogen, and ∼135 cleavage sites for MMP-2 to degrade collagen IV. For silk scaffolds formed with a mass ratio of silk to ECM molecule of 150:1, there will be negligible effects in *in vitro* degradation.

Additionally, *in vivo* heparin can bind to serine protease antithrombin III (AT).^85^ Upon binding, AT can undergo a confirmational change to prevent the activation of blood clotting proteases including serine proteases, thrombin, and factor Xa.^86-87^ So, if the heparin in the silk scaffold is not locked into the silk β-sheet structures, the addition of heparin could bind AT and change the rate of degradation.

### 2.3 Resolving differences in *in vivo* and *in vitro* results

Previous investigations with other biopolymers have often shown that *in vitro* and *in vivo* degradation can be divergent and that *in vitro* cell function and phenotypes within 3D materials do not always match *in vivo* results.^88-92^ We show using *in vitro* experimental data that addition of biopolymers to silk scaffolds does not change degradation (**Figure 4a-c, Table 2**). This is in direct contrast to our *in vivo* results which show that addition of biopolymers to silk scaffolds increases the rate of degradation *in vivo* (**Figure 3a, c, e, g**). This contrast of *in vitro* data and observations *in vivo* are critical for our understanding of the mechanism behind composite silk scaffold degradation. *In vitro* results demonstrate that enzyme-scaffold interactions are not the driving force for the data presented in **Figure 3a, c, e, g**. Instead, these results suggest that the types of cells infiltrating into the scaffold and the persistence of these cells within the scaffold as a function of bioactive molecule addition drive scaffold degradation.

#### 2.3.1 Scaffold composition dictates the infiltration and persistence of monocyte lineage cells in scaffolds over 8 weeks *in vivo*

**Figure 5a** shows ECM deposition (blue) at the interface between the scaffold and tissue in 5 scaffold formulations (silk, s-c, s-h, s-h-VEGF, s-c-h-VEGF) at 2 weeks and 8 weeks using Masson’s Trichrome staining. By observations of week 2 sections from all the conditions, silk scaffolds and scaffold compositions with VEGF had higher collagen deposition than s-c and s-h scaffolds (**Figure 5a1**). By week 8, observational assessment of all scaffolds suggests similar collagen deposition levels apart from s-h where most of the scaffold had degraded, and the area surrounding the remaining scaffold was populated with adipocytes (**Figure 5a2**).

**Figure 5.**
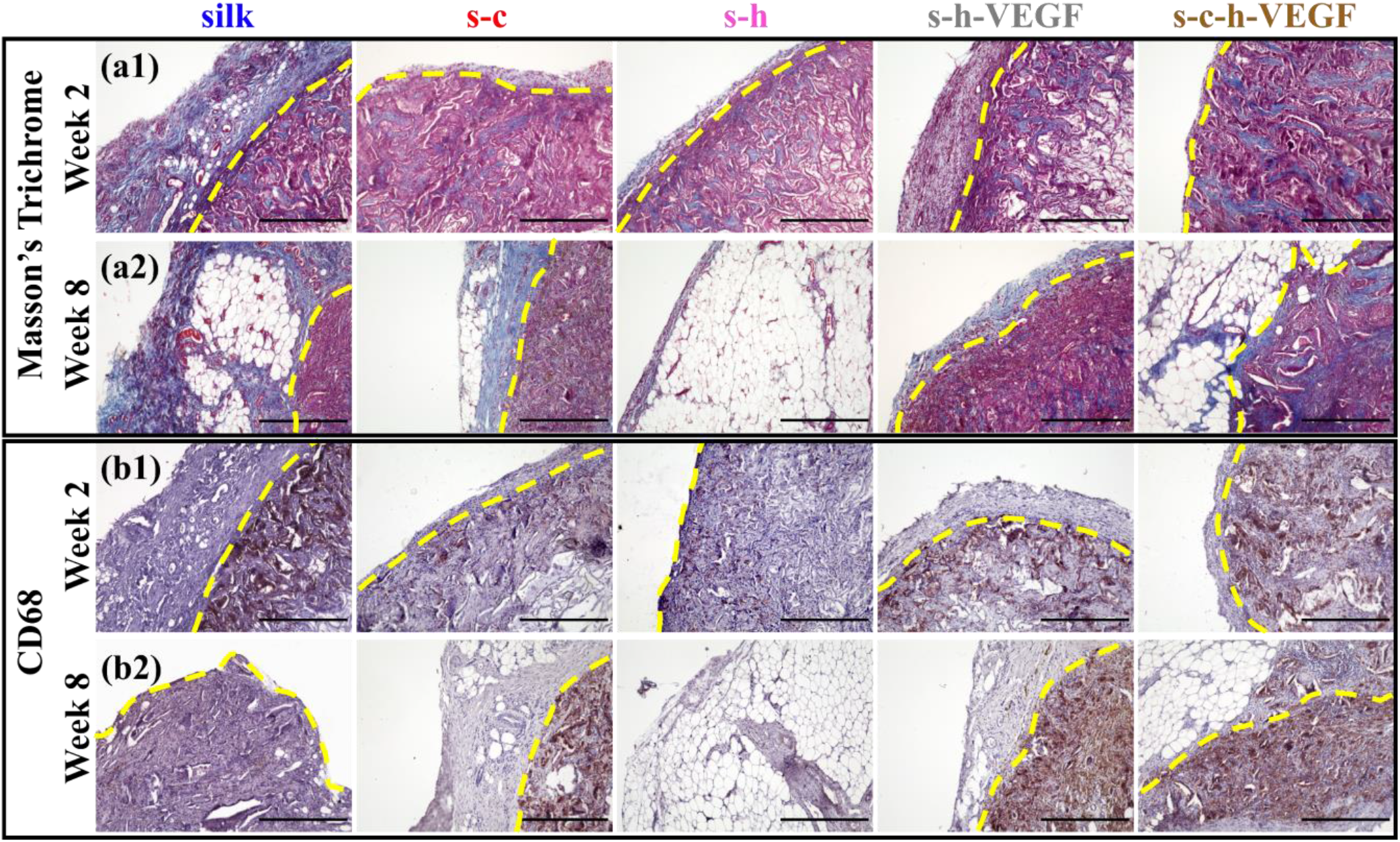
Scaffolds explanted 2 weeks (a1, b1) and 8 weeks (a2, b2) post-surgery were stained with (a) Masson’s Trichrome and (b) CD68+ cell (brown). Scale bars = 500 µm. The yellow dotted lines separate the scaffold and the overlaying excised tissue. The yellow dotted lines correspond to the black outline in Figure 2.

We aimed to further interrogate the types of cells infiltrating the scaffold, focusing on the role of monocyte-derived cells through detection of CD68. The presence of CD68 in and around the scaffolds at 2 and 8 weeks was evaluated with primary antibody staining and detection. For the purposes of this paper, we will refer to cells expressing any level of CD68 as CD68+ cells as confirming a specific phenotype. From **Figure 5b1**, we see that silk, s-h-VEGF and s-c-h-VEGF showed similar expressions of CD68+ cells near the scaffold-tissue interface as compared to s-c and s-h scaffolds at week 2. The addition of VEGF to scaffolds by pre (-) fabrication techniques promoted infiltration CD68+ cells. Multinucleated giant cells (MNGCs) were also observed in scaffolds with high expression of CD68 (**Figure S13**). By week 8, all scaffolds still had expression of CD68 regardless of VEGF incorporation and initial formulation with the lowest amount of CD68+ cells being observed within silk scaffolds (**Figure 5b2**). Moreover, CD68+ cells were only found within the scaffold, and not around the scaffold or in deposited tissue (**Figure 5b** and **Figure S14** where full scaffolds stained with CD68 are shown).

Following injury or biomaterial implantation, monocyte linage cells, such as macrophages, are one of the predominant immune cell classes known to hone to the site, but their spatiotemporal function is dependent upon cues from the local environment.^93^ These immune cells secrete MMPs that degrade components of the ECM and silk fibroin *in vivo*. At week 2, expression of CD68+ cells, MNGCs, and silk scaffold area are positively correlated, meaning the scaffolds with high expression of both CD68+ cells and MNGCs also had high scaffold areas. This suggests that at week 2, silk, s-h-VEGF, and s-c-h-VEGF scaffolds had not yet degraded completely as MNGCs were still present. Additionally, this positive correlation is the first to suggest that MNGCs are a potential mediator for silk-based scaffold degradation. By week 8, scaffold area was inversely correlated with CD68+ expression and MNGCs presence. Silk scaffolds had the lowest amount of CD68+ cells and the highest scaffold area at week 8, again suggesting that CD68+ cells are critical for scaffold degradation.

In addition, high collagen deposition (**Figure 5a**) trended with high levels CD68+ cells, except for in the silk only scaffolds. This phenomenon has also been suggested in other biomaterial formats^94-95^ and it is hypothesized that macrophages regulate fibrogenesis^96-97^ and recruit myofibroblasts, fibroblasts, and other inflammatory cells.^16, 98^ We aimed to evaluate other markers for cell phenotype such as myofibroblast activation and vascularization within these same scaffolds.

#### 2.3.2 Vascularization is dependent on scaffold composition *in vivo*

Both H&E and Masson’s Trichrome staining showed new blood vessel formation (**Figure S1, S2**). We leveraged the versatility of alpha smooth muscle actin (αSMA) staining to both track potential myofibroblasts or other activated cell types as well as locate maturing vessel structures. As qualitatively observed in **Figure 6a1**, by week 2, αSMA was expressed by cells at the leading edge of the infiltrating cell population for all scaffold conditions. In comparison of **Figure S14** vs. **Figure S15**, αSMA+ cells have infiltrated deeper into the scaffold compared to CD68+ cells at week 2. By week 8, αSMA staining was localized to blood vessels for all conditions, suggesting that myofibroblasts and activated cells are no longer present (**Figure 6a2**).

**Figure 6.**
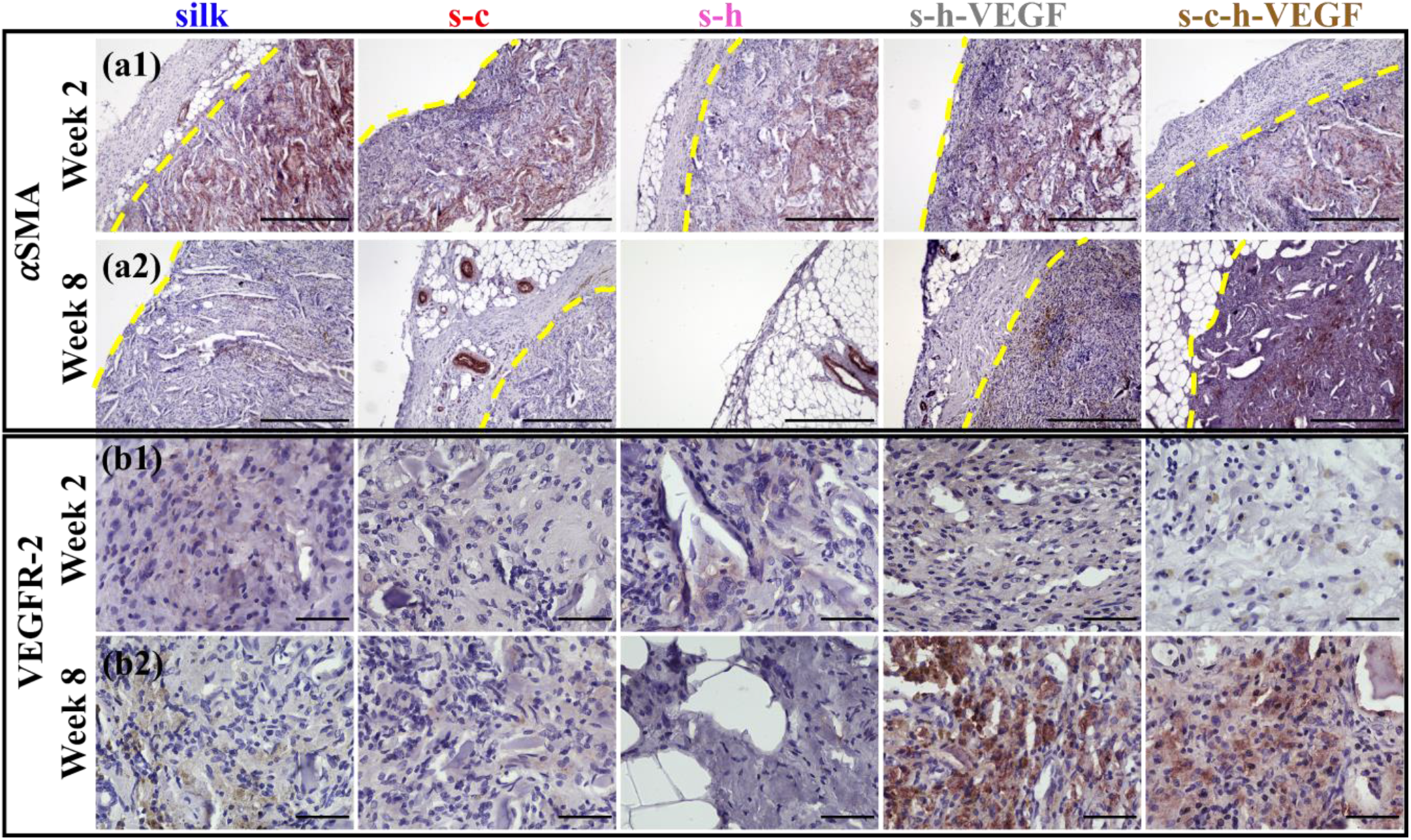
Scaffolds explanted 2 weeks (a1, b1) and 8 weeks (a2, b2) post-implantation were stained with (a) αSMA and (b) VEGFR-2. Scale bars (a) = 500 µm. Scale bars (b) = 50 µm. The yellow dotted lines separate the scaffold and the overlaying excised tissue. The yellow dotted lines correspond to the black outline in Figure 2.

To determine whether VEGF in the scaffold influences VEGFR-2 in the infiltrating cells to promote angiogenesis, we stained for VEGFR-2 at 2- and 8-weeks post-surgery (**Figure 6b**). At both week 2 and week 8, VEGFR-2 expression with scaffolds prefabricated with just collagen I or heparin was small compared to s-h-VEGF and s-c-h-VEGF scaffolds. The addition of VEGF to the scaffolds increases the expression of VEGFR-2 indicating the presence of endothelial cells.

VEGF loaded biomaterials are known to result in vascularization of new tissue formation.^99-101^ However, VEGF is expected to be short-lived after implantation and is prone to proteolytic degradation and low protein stability. In our study, the degradation of VEGF may have been slowed down by heparin immobilization. Our results show that VEGF initiated a sequence of events that underlies the prolonged VEGFR-2 expression over 8 weeks (**Figure 6b**). Instead of inducing potentially toxic responses to high systemic doses of VEGF, our system uses small concentration of VEGF immobilized to heparin for increased protection from proteolytic degradation and improved stimulation and proliferation of endothelial cells.

#### 2.3.3 Adipose tissue deposition is dependent on time post-surgery

To investigate the impacts of scaffold composition and fabrication methods on adipose tissue accumulation, we quantified the adipose area of each scaffold in both H&E- and Masson’s Trichrome-stained images (**Figure 7**). At early timepoints (week 1), we see that adipose tissue growth is mainly dependent on the scaffold fabrication method (**Figure 7a**). Scaffold formulation does not seem to have an impact early on in the adipocyte recruitment mechanism as evidenced by **Figure 7b-d**. At later timepoints, however, it becomes evident that adipose tissue growth is a function of time itself. At week 2, with the exception of scaffolds made without VEGF, adipose tissue deposition was significantly lower compared to silk for all fabrication methods and all formulations (**Figure 7a-d**). A similar trend is observed at week 8; for all formulations and for nearly all fabrication methods (excluding post (+,s) fabrication scaffolds), adipose tissue deposition was significantly greater in comparison with the silk condition (**Figure 7a-d**). Across all scaffold compositions and fabrication methods, there were no significant differences observed at week 4.

**Figure 7.**
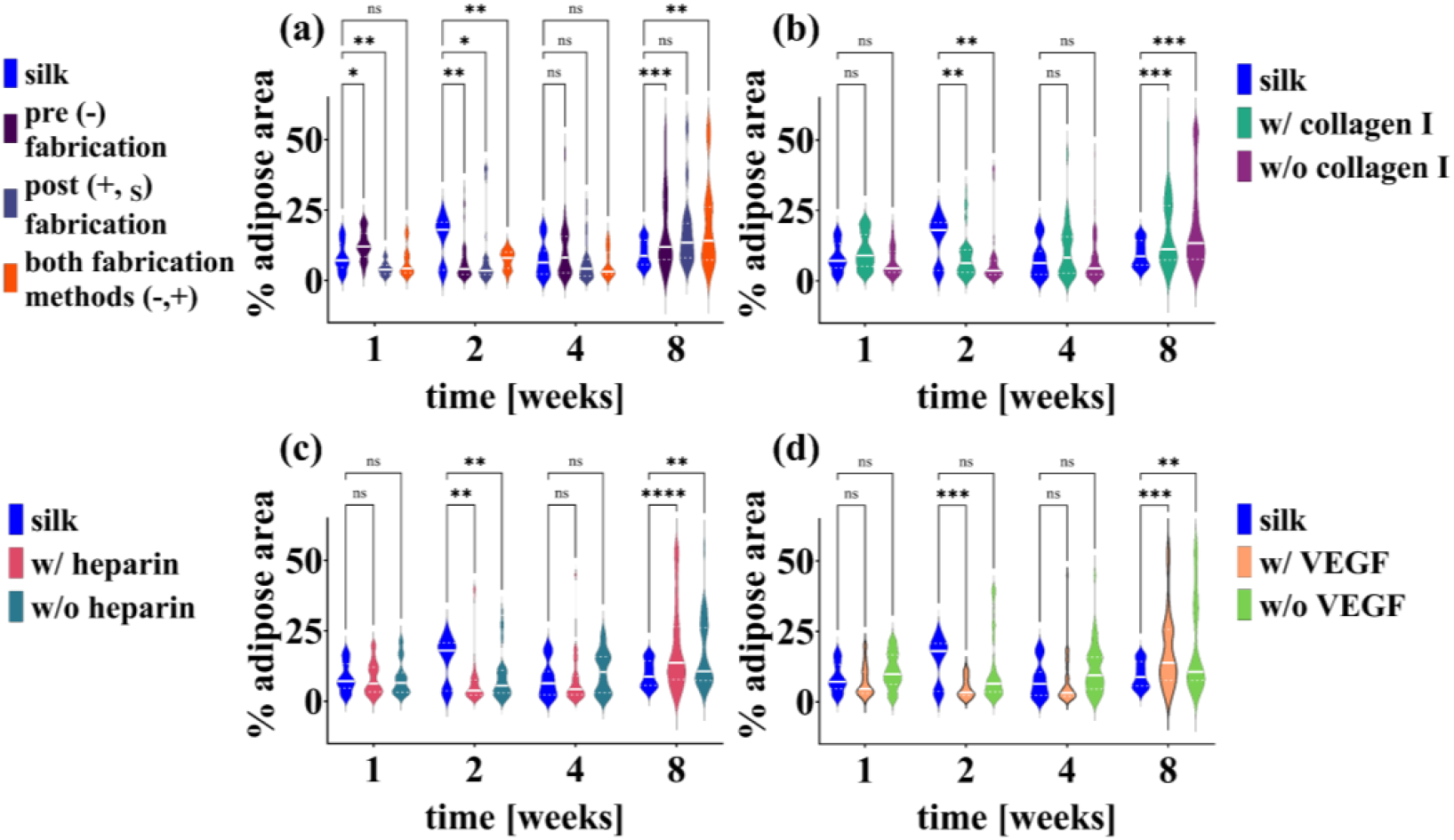
Quantitative changes of adipose area with scaffolds made from different fabrication conditions (a) and different bioactive molecules (b-d) after explant from a subcutaneous rodent model are presented here. Scaffolds were explanted 1-, 2-, 4-, and 8-weeks post-surgery. Statistical significance was determined using a mixed-effects model with Dunnett’s post-hoc multiple comparisons test. Statistical significance is reported as *p < 0.05, **p < 0.01, ***p < 0.001, and ****p < 0.0001.

For adipose tissue repair, investigators use composite materials with biological factors such as VEGF, FGF2, fibroblast growth factor 1 (FGF1), and insulin.^102-104^ The incorporation of these biological factors induce adipogenesis and vascularization. Vascularization in adipose tissue is critical to maintain access to oxygen and other nutrients. In one study, adipose tissue formation increased from 6 weeks to 12 weeks when bFGF was loaded using heparin in a decellularized scaffold.^102^ Adipogenesis takes 12 weeks indicating the presence of adipose tissue in and around the degrading scaffolds would likely increase after 8 weeks.^105^ The progressive adipose tissue deposition in our system is attributed to the slow release of VEGF or the prolonged effects initiated by VEGF.

## 3. Conclusions

Composite silk sponge-like scaffolds containing collagen I, heparin, and VEGF were used to systematically investigate growth factor delivery through modulation of enzymatic degradation. To explore a range of potential formulations to achieve tunable growth factor delivery, two fabrication methods (pre (-) fabrication and post (+, _S_) fabrication) were employed. Subcutaneous implants of composite silk scaffolds influenced recruitment of endogenous cells, scaffold degradation, local ECM deposition, and vascularization. To further characterize *in vivo* findings, we performed *in vitro* analyses. We determined that composite silk scaffolds have degradation rates dependent on local enzyme compositions. These results are the first step in building a promising pathway for personalized and tailored regenerative medicine. Future work aims to explore this system in injury models with defined functional recoveries as well as to investigate the use of this system with different growth factors.

## 4. Materials and Methods

### 4.1 Silk fibroin extraction

Through isolation from cocoons of *Bombyx mori* silkworms, silk fibroin solution was prepared as previously described.^106^ To isolate the pure silk fibroin protein, 5 g of silk cocoons were degummed in 2 L of boiling 0.02 M sodium carbonate (Catalog No. 451614, Sigma Aldrich, St. Louis, MO) in water for 30 minutes. Upon completion of degumming, the silk fibroin protein was air-dried in the fume hood for over 48 hours. To solubilize the dried fibroin protein, silk mats were denatured in 9.3 M aqueous lithium bromide (Catalog No. 213225, Sigma Aldrich, St. Louis, MO) at 60 °C for 4 hours. The solution was then dialyzed with a 3.5 kDa molecular weight cutoff membrane (Spectrum™ Spectra/Por™ 3 RC Dialysis Membrane Tubing, 3,500 Da MWCO, Catalog No. 08-670-5C, Thermo Fisher Scientific, Waltham, MA) to remove the lithium bromide ions in deionized water. To remove insoluble particles from the solubilized silk solution, the solubilized silk solution was centrifuged three times (>2000 g, 20 min, 4 °C). The concentration of the silk solution was determined by drying a known volume of the silk solution at 60 °C and massing the final dry weight following complete water loss. This protocol resulted in a 5–7% wt. v^-1^ (wt./v, 0.05-0.07 g/mL) silk solution. For a maximum of 3 weeks, silk solutions were stored at 4 °C prior to use in making scaffolds.

### 4.2 Collagen isolation

All animal experiments were executed under protocols approved by the Institutional Animal Care and Use Committees at University of Florida or Tufts University. These protocols are in accordance with the Guide for the Care and Use of Laboratory Animals (NIH, Bethesda MD). Ready-to-use reconstituted type I collagen was prepared as previously described.^107^ Briefly, from the tails of adult Sprague Dawley rats, rat tail type I collagen was isolated which will be referred to as collagen I in the manuscript. To obtain sterile soluble collagen I, the extracted collagen I was processed in acetic acid solution. Then the collagen I was frozen at -20 °C and lyophilized at -80 °C (FreeZone 12 Liter -84 °C Console Freeze Dryer, Labconco, Kansas City, MO). After lyophilization, the resulting collagen sponge-like material was solubilized in 0.02 N acetic acid.

### 4.3 Pre (-) fabrication scaffold formation

For incorporation into silk scaffolds, heparin (Heparin sodium salt from porcine intestinal mucosa, 250KU, Catalog No. H3149-250KU, Sigma Aldrich, St. Louis, MO, 100 µg/mL), collagen I (200 µg/mL), and/or VEGF (Catalog No. 100-20 Recombinant Human VEGF165, Peprotech, Rocky Hill, NJ, 0.1 µg/mL) were mixed before being added to diluted aqueous silk solution (final concentration of 3% wt. v^-1^ 0.03 g/mL). Isotropic scaffolds were formed by pouring the silk solution into wells of a 6-well plate and freezing the solution in a -80 °C freezer overnight. Following freezing, the frozen blocks were lyophilized at -80 °C (FreeZone 12 Liter -84 °C Console Freeze Dryer, Labconco, Kansas City, MO) for 5-7 days. The sponge-like scaffolds were partially crystalized by water annealing at room temperature under -0.05 MPa vacuum pressure with 500 mL of ultrapure Milli-Q water for 12 hours. The vacuum desiccator (Bel-Art SP™ Scienceware™ Lab Companion Round Style Vacuum Desiccator, Catalog No. 08-648-100, Thermo Fisher Scientific, Waltham, MA) has a volume of 6L. Water annealing induces β-sheet formation in the silk fibroin, resulting in silk protein crystallization.^25^ For *in vivo* experiments, sponge-like scaffolds were cut to a radius of 3 mm and height of 3 mm. For *in vitro* experiments, scaffolds were cut to a radius of 3 mm and height of 4 mm. Pre (-) fabrication scaffold formation is denoted by “-” in sample naming (**Figure 1, Table 1**). In all cases, scaffold architecture was inspected in each scaffold prior to use and if the porosity was non-uniform, that scaffold was not used in any experiments.

### 4.4 Post (+, _S_) fabrication scaffold formation

Scaffolds were prepared according to previous methods,^25^ following the steps identified above. However, bioactive molecules were not introduced to the scaffold until after it was partially crystallized, and the silk fibroin protein was no longer fully water soluble. VEGF and heparin were solubilized in 1X phosphate buffered saline (PBS), where subscript _S_ in VEGF_S_ and heparin_S_ designates soluble VEGF and heparin added post (+, _S_) fabrication, respectively. Scaffolds were soaked in solubilized solutions of VEGF_S_ and/or heparin_S_ for 30 minutes. Post (+, _S_) fabrication scaffold formation is denoted by “+” and “_S_” in sample naming (**Figure 1, Table 1**). In all cases, scaffold architecture was inspected in each scaffold prior to use and if the porosity was non-uniform, that scaffold was not used in any experiments.

### 4.5 *In vitro* scaffold evaluation

#### 4.5.1 Enzymatic degradation of scaffolds – Continuous Method

To experimentally determine the rate of scaffold degradation *in vitro* over time, scaffolds were kept in a well-mixed enzyme solution and the mass of the remaining scaffold was determined at specific time points as previously described.^26^ Briefly, the mass of dry scaffolds was recorded, and the scaffolds were placed in 1.5 mL Eppendorf tubes with 1 mL of proteinase K (1 U/mL and 0.1 U/mL) in PBS. Proteinase K (proteinase K from Tritirachium album lyophilized powder, Lot # 118M4089V, Catalog No. P6556, Sigma Aldrich, St. Louis, MO) had an activity of 30 U/mg. The Eppendorf tubes were incubated with continuous shaking at 37 °C and enzyme solution was removed and replaced every 48 hours. On days 1-7, 11, and 14scaffold dry mass was recorded. Prior to weighing, liquid was removed by first squeezing the sponge-like scaffold followed by aspiration of any remaining enzyme solution, resulting in a dry paper-like sample that rehydrated to original volume after submersion in solution again. Scaffold degradation was calculated as % remaining dry mass compared to original dry scaffold mass.

#### 4.5.2 Theoretical basis for the enzymatic degradation scaffolds *in vitro*

Lyophilized silk scaffold degradation has been previously modeled using two different kinetic models: Michaelis-Menten kinetics and modified first-order reaction kinetics.^26^ Each model is described in detail in previous work by Jameson and colleagues where the assumption is made that the dominant kinetic mode for the breakdown of the scaffold is through enzymatic degradation.^26^ It is important to emphasize that these models assume that the enzyme concentration is temporally and spatially uniform and well-mixed throughout the volume of the scaffold and surrounding bulk fluid. Likewise, the time-dependent concentration for the substrate is uniform throughout the scaffold. These assumptions are consistent with the design of the experiments, where the samples were under constant agitation and fresh enzyme solution was supplied every 48 hours to maintain constant activity. As the timescale for the degradation process is on the order of days, diffusion is assumed to be insignificant.

#### 4.5.3 Scanning electron microscopy (SEM)

To visualize scaffold morphology, scanning electron microscopy (SEM) was utilized. Scaffolds were allowed to air dry in the chemical hood for 4 days. Constructs were sliced to reveal internal scaffold surfaces. Dried samples were affixed to a special ZEISS/LEO SEM Pin Stub Mount with an adhesive sticker, Ø12.7mm x 9mm pin height (Catalog No. 16202, Ted Pella, Inc., Redding, CA), sputter-coated with a 5 nm thick gold layer and imaged with a FEI Nova NanoSEM 430 SEM at the University of Florida Nanoscale Research Facility.

### 4.6 *In vivo* scaffold evaluation

All procedures were conducted under animal protocols approved by the Institutional Animal Care and Use Committees at University of Florida or Tufts University. All animals used in this study were 7–8-week-old Sprague Dawley rats (∼225 grams, Charles River Laboratories, Wilmington, MA). After general anesthesia of oxygen and isoflurane, the back of the rat was shaved and the skin was prepped with betadine disinfectant, followed by alcohol wipe, and repeated three times. Up to three small longitudinal incisions were made through the skin of the rat and sterilized acellular silk scaffolds^27-28^ (3 mm radius x 3 mm height) were subcutaneously implanted in lateral pockets of each rat. The incisions were closed with surgical clips that were removed after 5-7 days. At week 1-, 2-, 4-, and 8-weeks post-surgery, animals were euthanized. The silk scaffolds along with the surrounding tissue (when adhered to the scaffold) were excised and collected for histological examination. The experimental design is described in **Table S2** where the rat numbers and conditions are described for each time point. Each scaffold composition was subcutaneously implanted with a biological replicate of at least n=3 for each time point.

#### 4.6.1 Histochemical analysis of *in vivo* samples

Samples were fixed with 10% phosphate buffered formalin (Catalog No. SF100, Thermo Fisher Scientific, Waltham, MA) and embedded in paraffin following a series of graded ethanol and xylene incubations. Samples were sectioned to 7 – 15 µm thickness and deparaffinized. Sections were either stained with Hematoxylin (Catalog No. SDGHS280, Thermo Fisher Scientific, Waltham, MA) and Eosin (Catalog No. SDHT1101128, Thermo Fisher Scientific, Waltham, MA) identified as H&E to visualize cell nuclei, Masson’s Trichrome (Catalog No. HT15, Sigma-Aldrich, St. Louis, MO) to visualize collagen deposition, or the following antibodies: alpha smooth muscle actin (αSMA, Catalog No. PA5-19465, Thermo Fisher Scientific, Waltham, MA), cluster of differentiation 68 (CD68, Catalog No. ab31630, Abcam, Cambridge, MA), and vascular endothelial growth factor receptor 2 (VEGFR-2, Catalog No. PA5-16487, Thermo Fisher Scientific, Waltham, MA). For sections being stained for a target antigen, heat-mediated antigen retrieval was performed using a citrate buffer. Secondary visualization of the primary antibodies was completed with ImmPRESS Horse anti-rabbit Ig Kit (Catalog No. MP-7401-50, Vector Laboratories, Burlingame, CA) or ImmPRESS Horse anti-mouse Ig Kit (Catalog No. MP-7422-15, Vector Laboratories, Burlingame, CA) and ImmPACT NovaRed substrate (Catalog No. SK-4805, Vector Laboratories, Burlingame, CA). In addition, these sections were then counterstained with Hematoxylin. Following staining and dehydration, samples were embedded in DPX Mountant (Catalog No. 50-980-370, Thermo Fisher Scientific, Waltham, MA) and imaged using a Keyence^®^ BZ-X800 series microscope.

#### 4.6.2 Scaffold area

To examine the degradation of scaffolds in an *in vivo* environment, histological images (H&E and Masson’s Trichrome) were used to quantify the scaffold area over time. Scaffold area (%) was calculated as

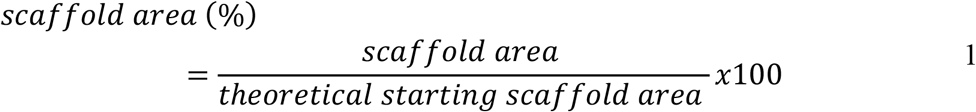

with a description of variables given in **Figure 2**. The theoretical starting scaffold area is calculated from *Area* =π*r*^2^ where *r* is the radius of the scaffold before implantation (*r* = 3 *mm*). At least five images from each time point, condition, and rat were quantified.

#### 4.6.3 Cell infiltration

Cell infiltration was quantified from H&E and Masson’s Trichrome stained sections. Cell infiltration area (%) was calculated as

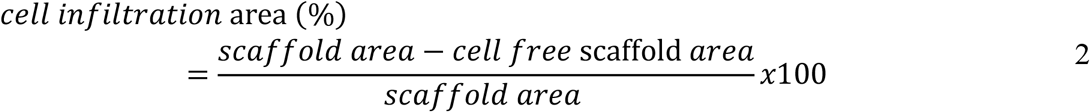

with a description of variables in **Figure 2**. At least five images from each time point, condition, and rat were analyzed.

#### 4.6.4 Adipose area

Adipose area was quantified from H&E and Masson’s Trichrome stained sections using a previously published convolutional neural network model named jointly optimized spatial histogram UNET architecture with an attention mechanism (JOSHUA+).^108^ Adipose area (%) was calculated as

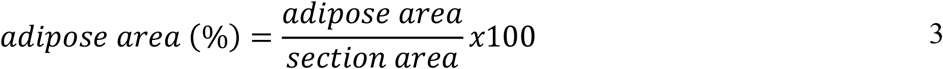

with a description of variables in **Figure 2**. At least five images from each time point, condition, and rat were analyzed.

### 4.7 Statistics

Experimental degradation data are expressed as mean ± 1 standard deviation (SD). Rate constant parameters were determined from weighted least squares analysis and standard deviation was calculated from Monte Carlo studies as previously reported.^26^

Quantification of scaffold area, cell infiltration area, and adipose area are expressed in violin plots. The middle white line is the median. The lower and upper white dotted lines are the 25^th^ and 75^th^ percentile, respectively. Mixed-effects model with the Geisser-Greenhouse correction was performed because equal variability of differences was not assumed. Dunnett’s test was used to correct for multiple comparisons. Statistical significance is reported as *p < 0.05, **p < 0.01, ***p < 0.001, and ****p < 0.0001.

## Supporting information

JamesonSupplementalMaterials

## 5. Acknowledgements and Funding

The authors would first like to acknowledge the efforts of undergraduate and graduate students that supported the final investigations presented here, including Kelly Bukovic, Alexa Espinoza and Keri Clarke, University of Florida Chemical Engineering Bachelors program graduates and Parth Patel, a University of Florida Chemical Engineering Master’s program graduate. The authors also thank the NIH (P41 EB002520) for support of this work in its initial stages. WLS acknowledges support from the National Institutes of Health Institutional Research and Academic Career Development Awards program at Tufts University (K12GM074869, Training in Education and Critical Research Skills (TEACRS)), which supported her prior to the start of her faculty position. JMG would like to acknowledge support from an NIH post-doctoral fellowship (F32-DE026058), which funded him prior to the start of his faculty position. JFJ acknowledges support from the University of Florida Herbert Wertheim College of Engineering Graduate School Preeminence Award and the University of Florida Institute for Cell and Tissue Science and Engineering Pittman Fellowship. MOP acknowledges support from the University of Florida Herbert Wertheim College of Engineering Graduate School Preeminence Award, the Kirkland Fellowship, and support from the Department of Defense (W81XWH2110199). We would also like to acknowledge the University of Florida Herbert Wertheim College of Engineering Research Service Center for help with data collection and analysis, especially Andres Trucco. Additionally, we would like to acknowledge Dr. Jason E. Butler for confirming our proper application of kinetic modeling^26^ and Dr. Alina Zare and Dr. Joshua Peeples for the collaborative work^108^ that lead to analysis of adipose tissue deposition as a function of scaffold formulation. The Stoppel Lab thanks the undergraduate students that helped with silk solution preparation and the initial startup of the lab at UF and the undergraduates that started the initial work at Tufts University. Parts of the Graphical Abstract and **Figure 1** were created with a license to BioRender.com.

