## Supplementary material for "Impact of bioactive molecule inclusion in lyophilized silk scaffolds varies between *in vivo* and *in vitro* assessments": JamesonSupplementalMaterials

**Additional information:**

**<u>Co-Author contact information:</u>**
Julie F. Jameson:
Marisa O. Pacheco:
Elizabeth C. Bender:
Nisha M. Kotta:
Lauren D. Black, III:
David L. Kaplan:
Jonathan M. Grasman:

### 1. Terminology used within this manuscript

Scaffold terminology used throughout the manuscript is given in **Table S1**. The colors, shapes, and naming are used in graphs and the main text to help the reader distinguish between sample types.

**Table S1. Scaffold terminology, scaffold abbreviation, and scaffold associated symbol for *in vitro* and *in vivo* investigations.**

| scaffold | abbreviation | symbol |
|---|---|---|
| silk | silk | ● |
| silk-collagen I | s-c | ■ |
| silk-heparin | s-h | ▲ |
| silk-collagen I-heparin | s-c-h | ◆ |
| silk-VEGF | s-VEGF | ⬢ |
| silk-collagen I-VEGF | s-c-VEGF | ⬟ |
| silk-heparin-VEGF | s-h-VEGF | ⬘ |
| silk-collagen I-heparin-VEGF | s-c-h-VEGF | ★ |
| silk+$\text{VEGF}_S$ | s+$\text{VEGF}_S$ | ▼ |
| silk+$\text{heparin}_S$ | s+$h_S$ | ⬒ |
| silk+$\text{heparin}_S$+$\text{VEGF}_S$ | s+$h_S$+$\text{VEGF}_S$ | ◓ |
| silk-collagen I+$\text{VEGF}_S$ | s-c+$\text{VEGF}_S$ | ✖ |
| silk-heparin+$\text{VEGF}_S$ | s-h+$\text{VEGF}_S$ | ✚ |

### 2. Representative histological images for each sample evaluated *in vivo*

**Figure S1, Figure S2 and Figure S3** show hematoxylin and eosin (H&E) staining of scaffolds recovered following 1-, 2-, 4-, and 8-weeks of implantation in a subcutaneous pocket of a 7-week-old Sprague Dawley rat. **Figure S4, Figure S5,** and **Figure S6** shows the same sets of samples stained with Masson's Trichrome. These images, and their counterparts from other samples, were evaluated following the outlines given in **Figure 2** of the manuscript to arrive at the data analyzed for cell infiltration and scaffold area. These images were also used in the calculation of adipose tissue using JOSHUA+.[1]

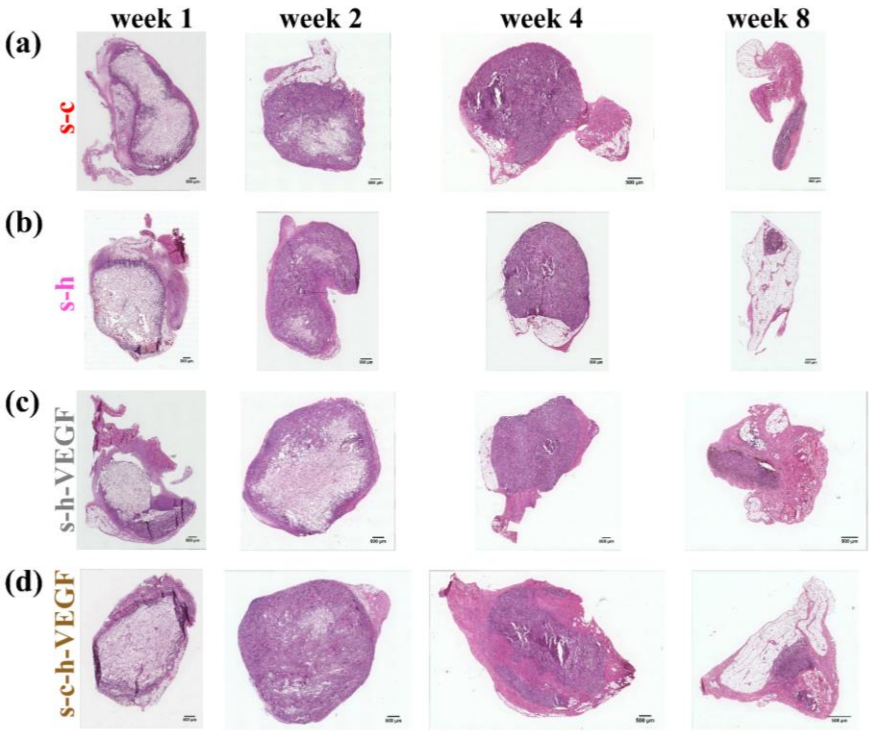


**Figure S1.** Representative H&E-stained images of explanted scaffolds at 1-, 2-, 4-, and 8-weeks post-surgery for pre (-) fabrication formulations: (a) silk-collagen I, (b) silk-heparin, (c) silk-heparin-VEGF, and (d) silk-collagen I-heparin-VEGF.

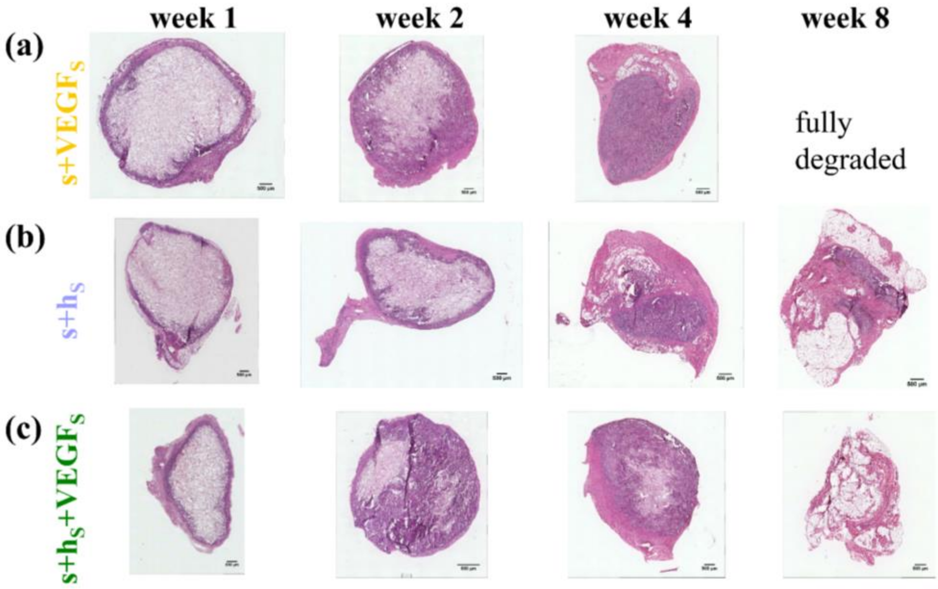


**Figure S2.** Representative H&E-stained images of explanted scaffolds at 1-, 2-, 4-, and 8-weeks post-surgery for post (+,$_S$) fabrication formulations: (a) silk+VEGF$_S$, (b) silk+heparin$_S$, and (c) silk+heparin$_S$+VEGF$_S$.

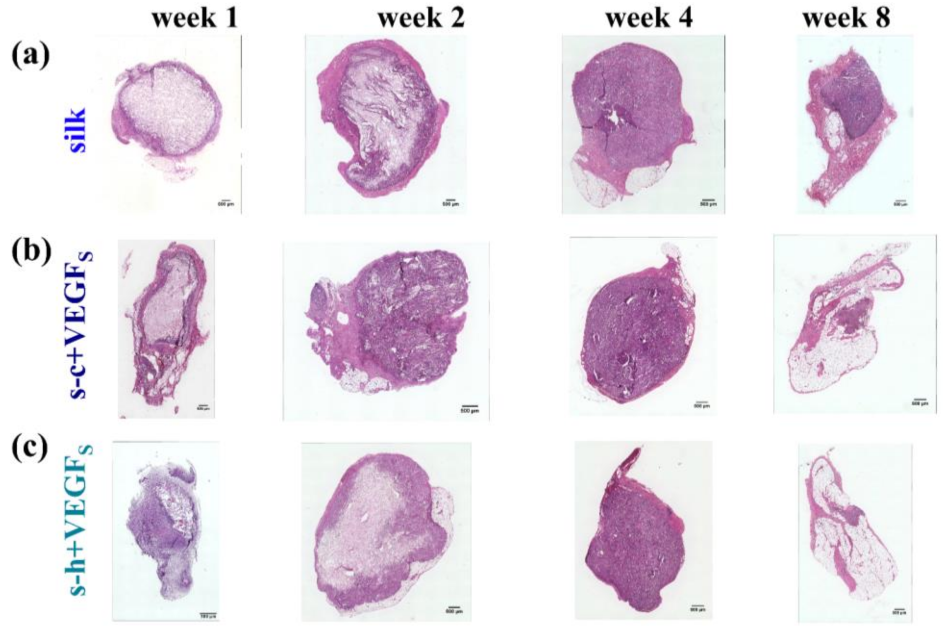


**Figure S3.** Representative H&E-stained images of explanted scaffolds at 1-, 2-, 4-, and 8-weeks post-surgery for post (+,$_S$) fabrication formulations.

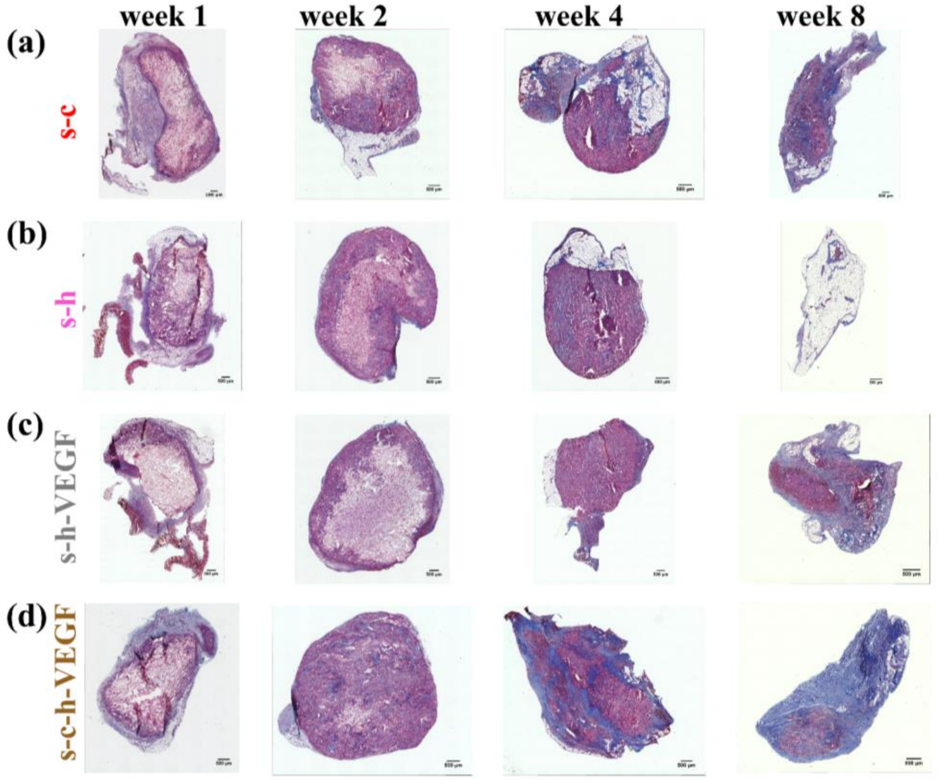


**Figure S4.** Representative H&E-stained images of explanted scaffolds at 1-, 2-, 4-, and 8-weeks post-surgery for (a) silk scaffolds and scaffolds formed with both (+,$_S$) fabrication methods: (b) silk-collagen+VEGF$_S$ and (c) silk-heparin+VEGF$_S$.

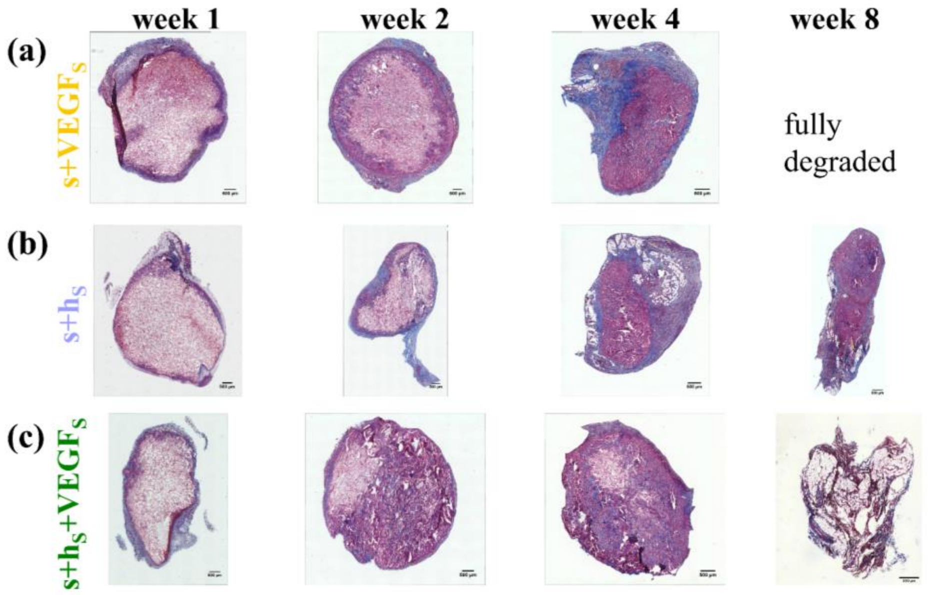


**Figure S5.** Representative Masson's Trichrome-stained images of explanted scaffolds at 1-, 2-, 4-, and 8-weeks post-surgery for pre (-) fabrication formulations: (a) silk-collagen I, (b) silk-heparin, (c) silk-heparin-VEGF, and (d) silk-collagen I-heparin-VEGF.

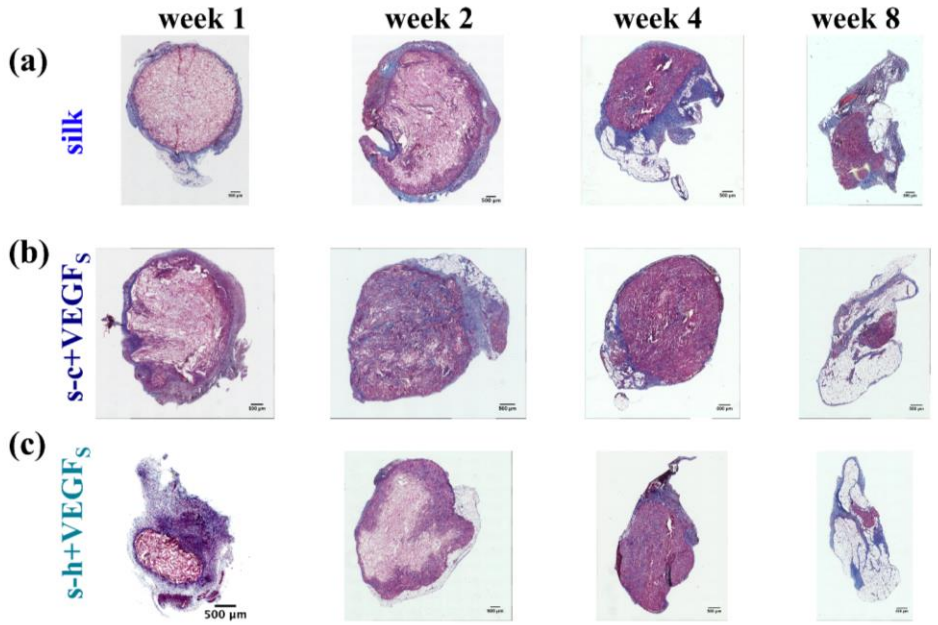


**Figure S6.** Representative Masson's Trichrome-stained images of explanted scaffolds at 1-, 2-, 4-, and 8-weeks post-surgery for post (+,$_S$) fabrication formulations: (a) silk+VEGF$_S$, (b) silk+heparin$_S$, and (c) silk+heparin$_S$+VEGF$_S$.

### 3. Statistical comparisons for individual formulations after *in vivo* image analysis and calculation

While the many results in the main text discuss combined formulations to assess the individual impacts of heparin, collagen I, and VEGF inclusion and also assess the method of scaffold preparation, individual comparisons are provided in this supplement. The results presented follow the determination of significance using a mixed-effects model with Šídák's post-hoc multiple comparisons test. Statistical significance is reported as *p < 0.05, **p < 0.01, ***p < 0.001, and ****p < 0.0001 for all data in this section. **Figure S7** compares total remaining scaffold area for scaffolds containing collagen I and the respective formulation control. For example, silk only scaffolds are compared to silk scaffolds containing collagen I in **Figure S7a**, while the addition of collagen I along with the use of soluble VEGF is shown in **Figure S7b**. Finally, **Figure S7c** shows a comparison between the addition of collagen for scaffolds formed with both heparin and VEGF. **Figure S8** shows a similar set of comparisons for heparin-containing scaffolds while **Figure S9** shows comparisons for scaffolds with VEGF. Similarly, cell infiltration comparisons are given in **Figure S10**, **Figure S11**, and **Figure S12**.

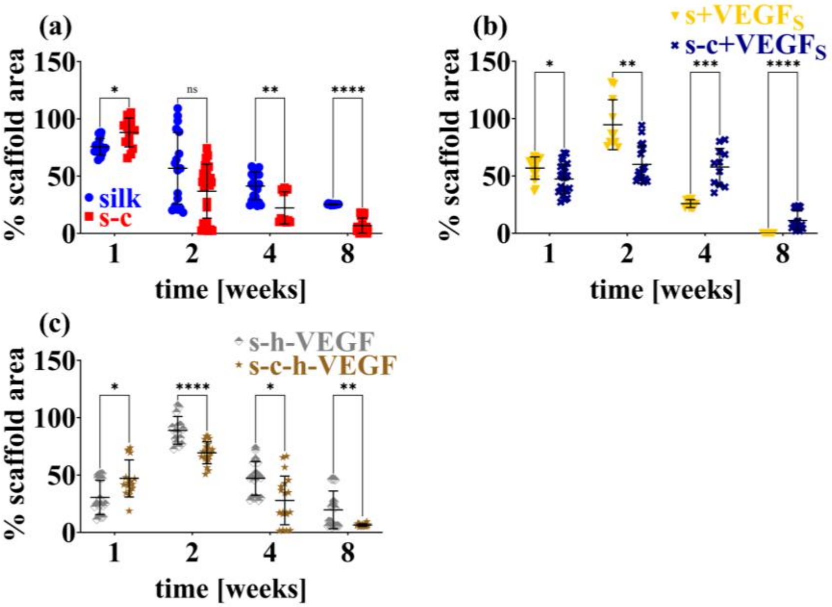


**Figure S7**. Scaffold area percent for negative (scaffolds without collagen I) and positive (scaffolds with collagen I) controls over 8 weeks. (a) Silk and silk-collagen I scaffold areas over 8 weeks. (b) Silk+VEGF$_S$ and silk-collagen I+VEGF$_S$ scaffold areas over 8 weeks. (c) Silk-heparin-VEGF and silk-collagen I-heparin-VEGF scaffold areas over 8 weeks. Statistical significance was determined using a mixed-effects model with Šídák's post-hoc multiple comparisons test. Statistical significance is reported as *p < 0.05, **p < 0.01, ***p < 0.001, and ****p < 0.0001.

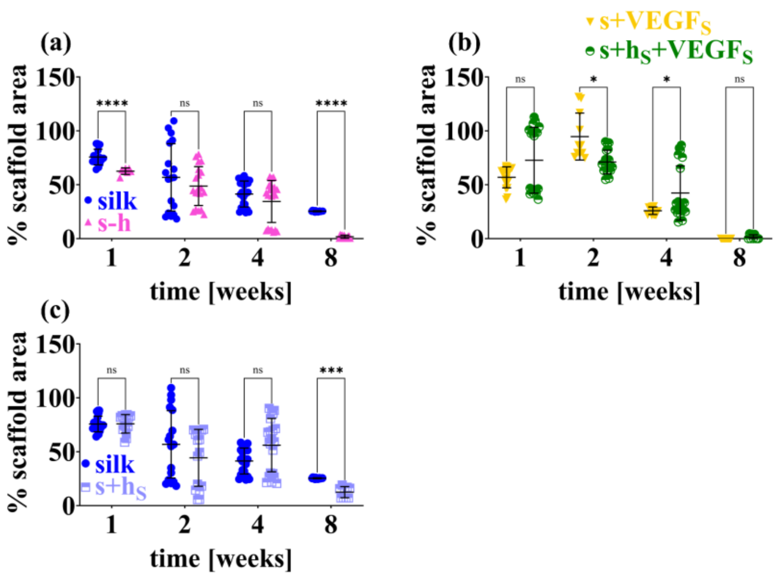


**Figure S8.** Scaffold area percent for negative (scaffolds without heparin) and positive (scaffolds with heparin) controls over 8 weeks. (a) Silk and silk-heparin scaffold areas over 8 weeks. (b) Silk+VEGF$_S$ and silk+heparin$_S$+VEGF$_S$ scaffold areas over 8 weeks. (c) Silk and silk+heparin$_S$ scaffold areas over 8 weeks. Statistical significance was determined using a mixed-effects model with Šídák's post-hoc multiple comparisons test. Statistical significance is reported as *p < 0.05, **p < 0.01, ***p < 0.001, and ****p < 0.0001.

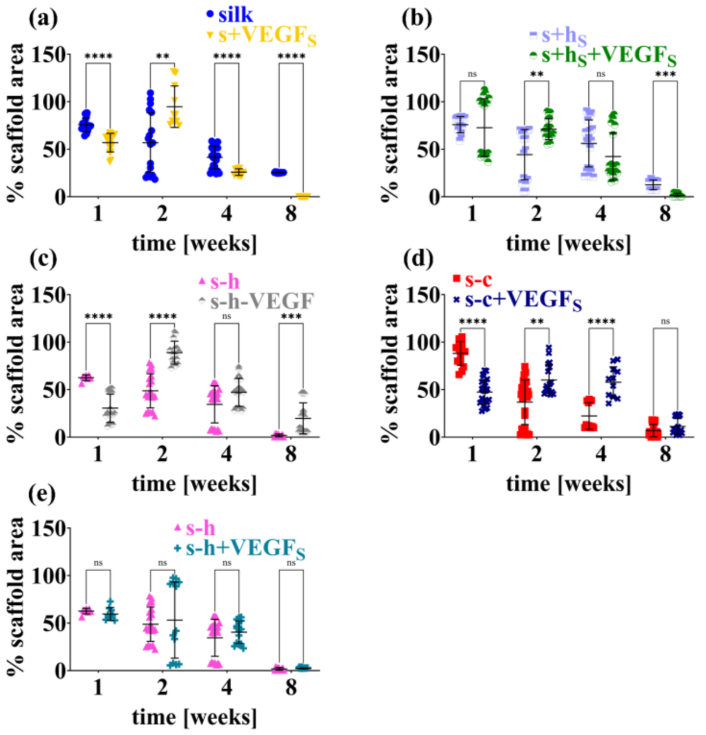


**Figure S9.** Scaffold area percent for negative (scaffolds without VEGF) and positive (scaffolds with VEGF) controls over 8 weeks. (a) Silk and silk+VEGF$_S$ scaffold areas over 8 weeks. (b) Silk+heparin$_S$ and silk+heparin$_S$+VEGF$_S$ scaffold areas over 8 weeks. (c) Silk-heparin and silk-heparin-VEGF scaffold areas over 8 weeks. (d) Silk-collagen I and silk-collagen I+VEGF$_S$ scaffold areas over 8 weeks. (e) Silk-heparin and silk-heparin+VEGF$_S$ scaffold areas over 8 weeks. Statistical significance was determined using a mixed-effects model with Šídák's post-hoc multiple comparisons test. Statistical significance is reported as *p < 0.05, **p < 0.01, ***p < 0.001, and ****p < 0.0001.

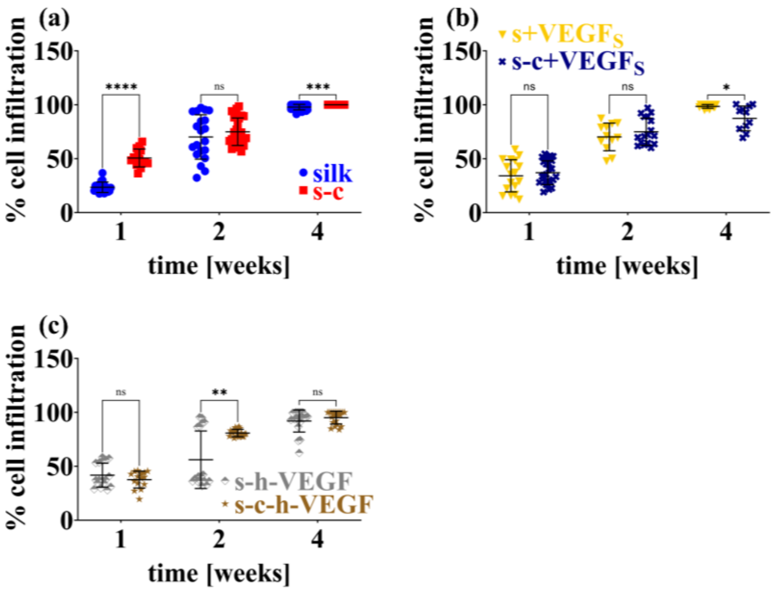


**Figure S10.** Cell infiltration percent for negative (scaffolds without collagen I) and positive (scaffolds with collagen I) controls over 8 weeks. (a) Silk and silk-collagen I cell infiltration areas over 8 weeks. (b) Silk+VEGF$_S$ and silk-collagen I+VEGF$_S$ cell infiltration areas over 8 weeks. (c) Silk-heparin-VEGF and silk-collagen I-heparin-VEGF cell infiltration areas over 8 weeks. Statistical significance was determined using a mixed-effects model with Šídák's post-hoc multiple comparisons test. Statistical significance is reported as *$p < 0.05$, **$p < 0.01$, ***$p < 0.001$, and ****$p < 0.0001$.

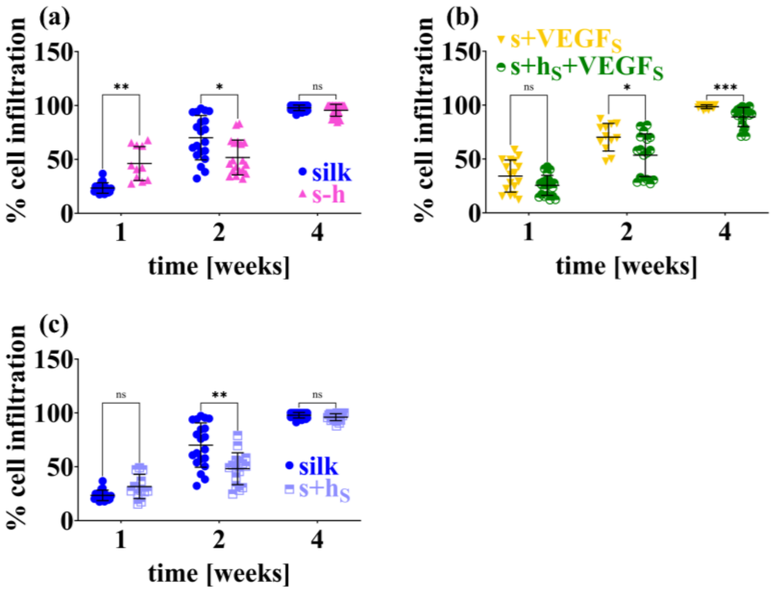


**Figure S11.** Cell infiltration percent for negative (scaffolds without heparin) and positive (scaffolds with heparin) controls over 8 weeks. (a) Silk and silk-heparin cell infiltration areas over 8 weeks. (b) Silk+VEGF$_S$ and silk+heparin$_S$+VEGF$_S$ cell infiltration areas over 8 weeks. (c) Silk and silk+heparin$_S$ cell infiltration areas over 8 weeks. Statistical significance was determined using a mixed-effects model with Šídák's post-hoc multiple comparisons test. Statistical significance is reported as *$p < 0.05$, **$p < 0.01$, ***$p < 0.001$, and ****$p < 0.0001$.

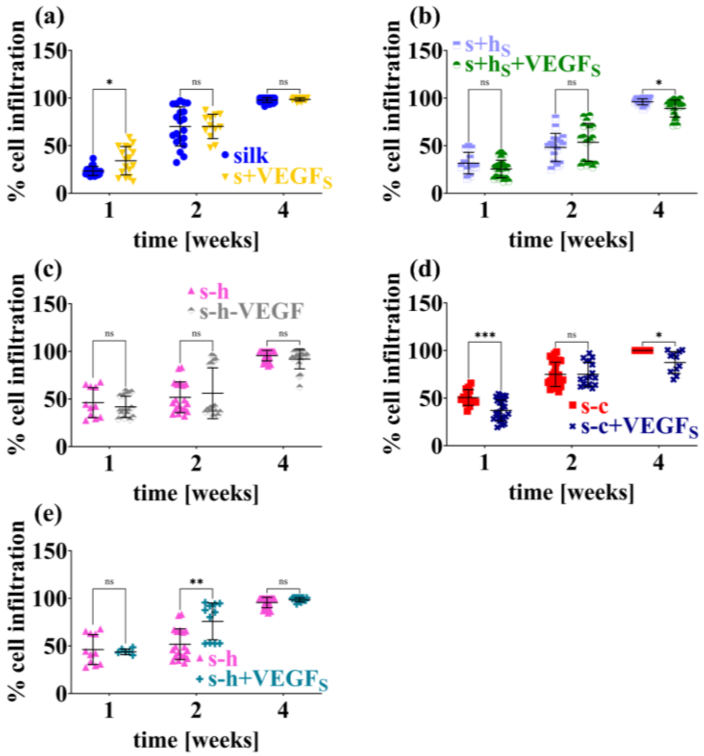


**Figure S12.** Cell infiltration percent for negative (scaffolds without VEGF) and positive (scaffolds with VEGF) controls over 8 weeks. (a) Silk and silk+VEGF$_S$ cell infiltration areas over 8 weeks. (b) Silk+heparin$_S$ and silk+heparin$_S$+VEGF$_S$ cell infiltration areas over 8 weeks. (c) Silk-heparin and silk-heparin-VEGF cell infiltration areas over 8 weeks. (d) Silk-collagen I and silk-collagen I+VEGF$_S$ cell infiltration areas over 8 weeks. (e) Silk-heparin and silk-heparin+VEGF$_S$ cell infiltration areas over 8 weeks. Statistical significance was determined using a mixed-effects model with Šídák's post-hoc multiple comparisons test. Statistical significance is reported as *$p < 0.05$, **$p < 0.01$, ***$p < 0.001$, and ****$p < 0.0001$.

### 4. Immunohistochemical evaluation of antibody expression as a function of bioactive molecule incorporation

Zoomed in image sections (**Figure S13**) and additional full section images (**Figure S14, Figure S15**) for the data presented in the manuscript in **Figures 5 and 6** are given in **Figure S13, Figure S14**, and **Figure S15** for CD68 and alpha smooth muscle actin (αSMA) to aid in the reader's interpretation of the trends observed and discussed in the text.

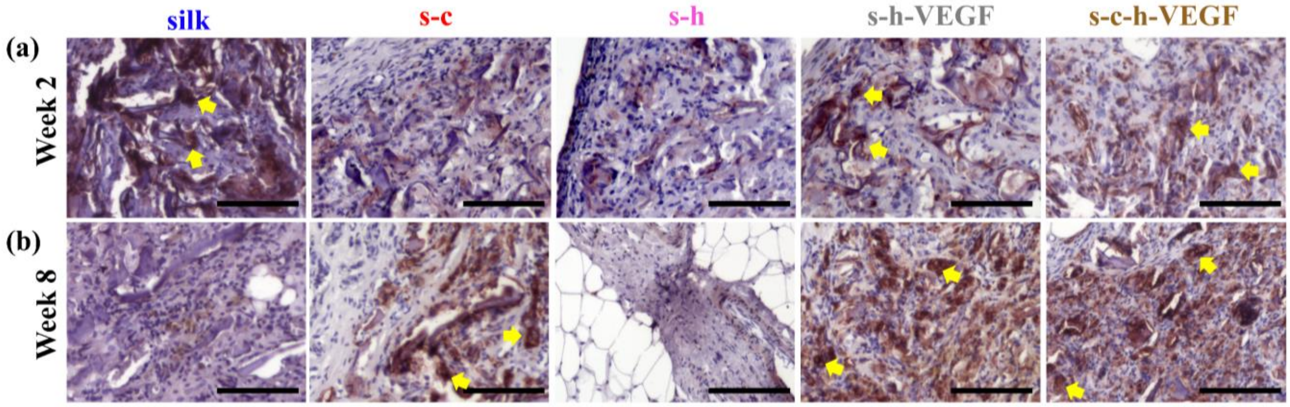


**Figure S13.** Immunohistochemical detection of CD68 (brown) in explanted scaffolds formed using the pre (-) fabrication method (a) 2 weeks and (b) 8 weeks post-surgery. Scale bars = 125 μm. The yellow arrows point toward multinucleated giant cells (MNGCs).

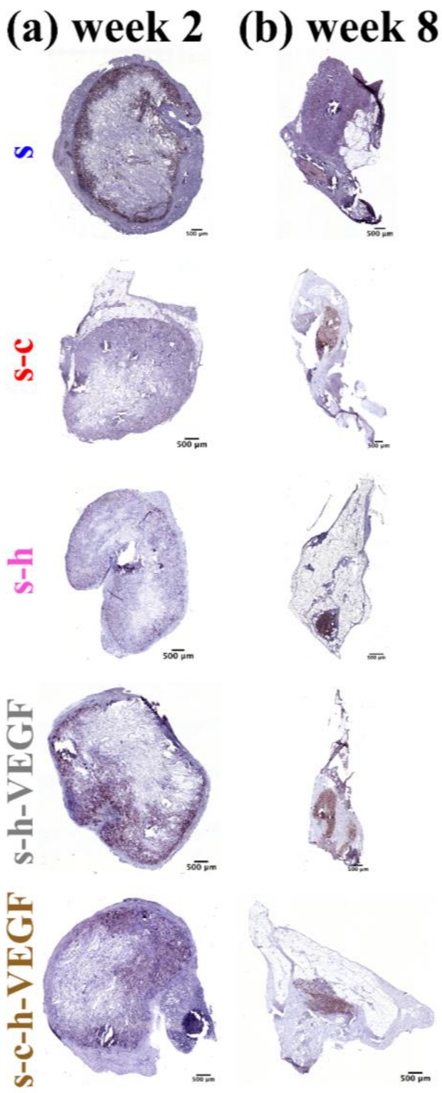


**Figure S14.** Immunohistochemical detection of CD68 (brown) in explanted scaffolds formed using the pre (-) fabrication method (a) 2 weeks and (b) 8 weeks post-surgery. Scale bars = 500 μm.

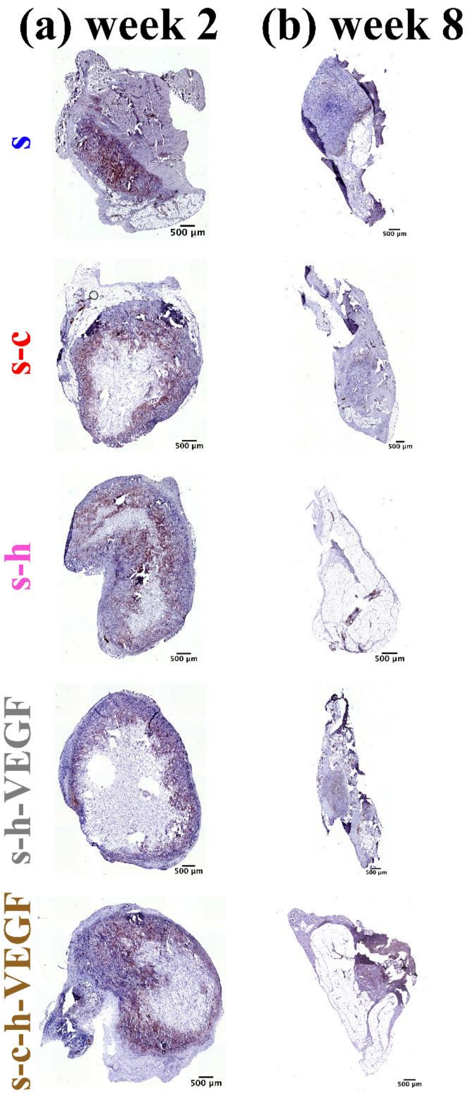


**Figure S15.** Immunohistochemical detection of αSMA (brown) in explanted scaffolds formed using the pre (-) fabrication method (a) 2 weeks and (b) 8 weeks post-surgery. Scale bars = 500 µm.

### 5. Subcutaneous implant study design and experimental details

**Table S2** provides the implant organization and study design for the *in vivo* studies. Implants were placed subcutaneously as described in the Materials and Methods section of the manuscript. For rats receiving eight scaffolds, 2 mm long incisions were made, and 2 scaffolds were placed along each side of the incision. For rats receiving ten scaffolds, 2 mm long incisions and 1 mm long incisions were made. One scaffold was placed on either side of the 1 mm incision while two were placed on each side for the 2 mm incision. At each time point, samples were recovered following humane euthanasia using $CO_2$ and cervical dislocation as approved by the Tufts University Institutional Animal Care and Use Committee (IACUC). After 8 weeks, some of the scaffolds were completely degraded and not recoverable.

**Table S2. Experimental design for *in vivo* subcutaneous model. All scaffolds were implanted on day 1 but harvested at later time points (columns). The scaffold formulation and its location relative to other formulations are represented by each square, as denoted by the asterisk(*) for scaffold implantation in the graphical abstract.**

| week 1 | | week 2 | | week 4 | | week 8 | |
|---|---|---|---|---|---|---|---|
| Rat 1 | | Rat 4 | | Rat 8 | | Rat 12 | |
| s | s-h+VEGF$_s$ | s | s-h | s | s-h | s | s-h |
| s+VEGF$_s$ | s-h-VEGF | s+VEGF$_s$ | s-h+VEGF$_s$ | s+VEGF$_s$ | s-h+VEGF$_s$ | s+VEGF$_s$ | s-h+VEGF$_s$ |
| s+h$_s$ | s-c | s+h$_s$ | s-h-VEGF | s+h$_s$ | s-h-VEGF | s+h$_s$ | s-h-VEGF |
| s+h$_s$+VEGF$_s$ | s-c+VEGF$_s$ | s+h$_s$+VEGF$_s$ | s-c | s+h$_s$+VEGF$_s$ | s-c | s+h$_s$+VEGF$_s$ | s-c |
| s-h | s-c-h-VEGF | | | | | | |
| Rat 2 | | Rat 5 | | Rat 9 | | Rat 13 | |
| s | s-c-h-VEGF | s-c+VEGF$_s$ | s-h | s-c+VEGF$_s$ | s-h | s-c+VEGF$_s$ | s-h |
| s+h$_s$ | s-c | s-c-h-VEGF | s-h-VEGF | s-c-h-VEGF | s-h-VEGF | s-c-h-VEGF | s-h-VEGF |
| s-h | s-h+VEGF$_s$ | s | s-c+VEGF$_s$ | s | s-c+VEGF$_s$ | s | s-c+VEGF$_s$ |
| s-h-VEGF | s+h$_s$+VEGF$_s$ | s+h$_s$ | s-c-h-VEGF | s+h$_s$ | s-c-h-VEGF | s+h$_s$ | s-c-h-VEGF |
| s-c+VEGF$_s$ | s+VEGF$_s$ | | | | | | |
| Rat 3 | | Rat 6 | | Rat 10 | | Rat 14 | |
| s-c-h-VEGF | s+h$_s$ | s-c | s-c-h-VEGF | s-c | s-c-h-VEGF | s-c | s-c-h-VEGF |
| s-h | s-h-VEGF | s-h+VEGF$_s$ | s-h | s-h+VEGF$_s$ | s-h | s-h+VEGF$_s$ | s-h |
| s-c+VEGF$_s$ | s+VEGF$_s$ | s+h$_s$+VEGF$_s$ | s-c+VEGF$_s$ | s+h$_s$+VEGF$_s$ | s-c+VEGF$_s$ | s+h$_s$+VEGF$_s$ | s-c+VEGF$_s$ |
| s+h$_s$+VEGF$_s$ | s-h+VEGF$_s$ | s+VEGF$_s$ | s+h$_s$+VEGF$_s$ | s+VEGF$_s$ | s+h$_s$+VEGF$_s$ | s+VEGF$_s$ | s+h$_s$+VEGF$_s$ |
| s-c | s | | | | | | |
| | | Rat 7 | | Rat 11 | | Rat 15 | |
| | | s-c | s-h+VEGF$_s$ | s-c | s-h+VEGF$_s$ | s-c | s-h+VEGF$_s$ |
| | | s+h$_s$ | s | s+h$_s$ | s | s+h$_s$ | s |
| | | s-h-VEGF | s-c | s-h-VEGF | s-h | s-h-VEGF | s-c-h-VEGF |
| | | s+VEGF$_s$ | s-c-h-VEGF | s+VEGF$_s$ | s | s+VEGF$_s$ | s-h-VEGF |
